# Livestock Landscapes as Ecological Filters: Effects of the Tree Cover Gradient on the Taxonomic and Functional Diversity of Granivorous Birds in the Colombian Amazon

**DOI:** 10.1101/2025.07.29.667560

**Authors:** Jenniffer Tatiana Díaz-Cháux, Alexander Velasquez-Valencia, Alejandro Navarro-Morales, Fernando Casanoves

## Abstract

Livestock landscapes in the Colombian Amazon are undergoing rapid land-use changes, leading to the structural simplification of vegetation and the loss of avifauna and its ecological functions. This study evaluated the effect of tree cover as an ecological filter on the taxonomic and functional diversity of granivorous birds within livestock mosaics. Eight livestock landscape mosaics were analyzed and classified based on tree cover into open, semi-open, and semi-closed categories. Bird surveys were conducted using point counts and mist netting, and eight morphological traits were measured across the 22 recorded species. Five functional diversity indices were calculated to assess the response to the tree cover gradient through their morphological traits. A reduction in tree cover excludes functionally specialized species and favors generalist communities with redundant traits. Semi-closed and semi-open covers harbored higher functional and taxonomic diversity, showing high levels of richness and functional evenness, with niche complementarity. Open pastures exhibited lower trait variability and clustering with reduced ecological functionality. Larger granivorous species were associated with denser vegetation, whereas smaller species were linked to more open, highly disturbed areas. Greater tree cover promotes the coexistence of granivorous bird species by increasing the availability of microhabitats and resource diversity, which are exploited by species with different functional traits. This highlights the importance of conserving structurally complex vegetation within livestock matrices, using silvopastoral systems to enhance the resilience and functionality of productive Amazonian landscapes.

## Introduction

The Colombian Amazon represents one of the most dynamic and diverse socioecological systems on the American continent. However, in recent decades, it has suffered the loss of more than two million hectares of natural forests due to the expansion of the agricultural frontier, land grabbing, and the increase in areas cultivated with illicit crops [1–3]. These drivers of deforestation have led to a transformation in land use, shifting from landscapes with high tree cover heterogeneity to matrices dominated by pastures surrounding small patches of remnant forest [4,5].

In this context, cattle ranching has become the predominant productive activity in the Colombian Amazon, with an inventory exceeding three million head of cattle per year [6]. The production of a single head of cattle requires more than 1.4 hectares of pasture, highlighting the low productivity of Amazonian soils for livestock farming [7,8]. According to Velasquez-Valencia and Bonilla-Gomez [9], livestock landscapes are composed of mosaics with low heterogeneity and poor connectivity between patches of arboreal or transitional vegetation. The current configuration of this landscape results from the historical interaction between land-use change and local physiography [10]. Its composition is characterized by small patches of agricultural land, agroforestry crops such as cacao (*Theobroma cacao* L.) and rubber (*Hevea brasiliensis* Müll. Arg.) generally smaller than 2 hectares, and narrow ecological corridors of forest remnants associated with lotic and lentic water bodies, all embedded within a matrix of pastureland [11,12].

The ecological dynamics of livestock landscapes in the Amazon are shaped by the ongoing loss of tree cover, which affects the local persistence of species [13–15]. The response of biological communities to habitat disturbance varies according to the ecological and functional attributes of species, such as dispersal capacity, reproductive efficiency, and food availability [16–18].

According to Parajuli [19] and Melo et al. [20], functional ecology provides a conceptual framework to analyze how species and communities respond to disturbance gradients. These studies are based on the evaluation of morphological, physiological, and ecological traits. Consequently, the loss of species with specialized traits may reduce functional diversity and compromise essential ecosystem functions [21–23]. Birds are highly sensitive to changes in landscape structure and composition; thus, they are an ideal biological group for evaluating ecological processes, ecosystem functioning, and functional diversity [24–26]. The latter is understood as the variety and distribution of traits that determine species persistence and their contribution to ecosystem processes [27]. From this perspective, the impacts of agroecosystems on biodiversity can be more accurately assessed using functional indicators, with bird assemblages serving as effective models [28–30].

Birds fulfill key ecological roles such as seed dispersal, biological pest control, pollination, and nutrient cycling [31–33]. These functions are modulated by the diversity of functional traits they possess, which determine reproduction, diet, and habitat use across gradients of vegetation cover [34]. Thus, functional traits are considered predictors of ecological interactions and community responses to habitat transformation [35,36]. It has been observed that bird assemblages in agroecosystems exhibit lower functional diversity than those in secondary forests, suggesting processes of functional homogenization driven by anthropogenic disturbance [29]. According to Navarro et al. [22], granivorous birds, despite their ecological plasticity, are sensitive to landscape fragmentation, leading to the loss of species with specialized functional traits and a reduction in overall functional diversity. Therefore, exploring the relationship between habitat loss and functional diversity is essential to understand the mechanisms of community assembly and environmental filtering; furthermore, it can provide critical information for conservation planning under scenarios of climate change [37].

To quantify the functional responses of communities, several multidimensional indices of functional diversity have been proposed, including FRic, FEve, FDis, FDiv, and RaoQ [38–42]. These indices allow researchers to link biodiversity with ecosystem functioning through the functional traits of species [17,43,44]. The Functional Richness index (FRic) estimates the extent of functional space occupied by the community; Functional Evenness (FEve) evaluates the regularity of trait distribution; Functional Divergence (FDiv) indicates how far dominant species are from the functional centroid; Functional Dispersion (FDis) reflects the average distance of species to the centroid of the functional space, influenced by dominant species; and Rao’s Quadratic Entropy (RaoQ) integrates functional dissimilarity weighted by relative abundance [38,44,45]. In addition, the RLQ multivariate analysis enables the identification of co-structuring patterns between environmental variables, functional traits, and species composition, thereby detecting processes of environmental filtering or functional selection [46].

This study assessed how the gradient of tree cover influences the structure, composition, and functional diversity of granivorous bird communities in livestock landscapes of the Colombian Amazon. The central hypothesis proposed that the tree cover gradient acts as an ecological filter that shapes the taxonomic and functional diversity of these communities. It is predicted that denser vegetation covers will host species with specialized functional traits, while more open covers will favor generalist, small-bodied species adapted to disturbed environments. The findings of this research provide insight into general rules of functional community assembly and help elucidate how environmental filters influence the functional structure of bird communities in productive Amazonian landscapes.

## Materials and methods

### Study area

The research was conducted in the northwestern region of the Colombian Amazon, across eight livestock landscape mosaics located in seven municipalities within the department of Caquetá: (i) Albania; 1.302766° N – 75.909201° W, 277 m a.s.l., (ii) El Doncello; 1.643959° N – 75.261978° W, 359 m a.s.l., (iii) Florencia; 1.429608° N – 75.528399° W, 248 m a.s.l., (iv) La Montañita; 1.336173° N – 75.352393° W, 269 m a.s.l., (v) Milán; 1.300662° N – 75.519043° W, 222 m a.s.l., (vi) Morelia; 1.409152° N –75.727271° W, 259 m a.s.l., and (vii) San José del Fragua; 1.270573° N – 76.007303° W, 376 m a.s.l. (Fig 1). The landscapes of the region are characterized by a geomorphology of low hills and piedmont, with floodable valleys and an average slope of 12%. The area has an average temperature of 28.62 °C, relative humidity of 86%, and annual precipitation of 4277 mm, with a unimodal distribution and a peak rainy season from April to October [47].

**Fig 1.**
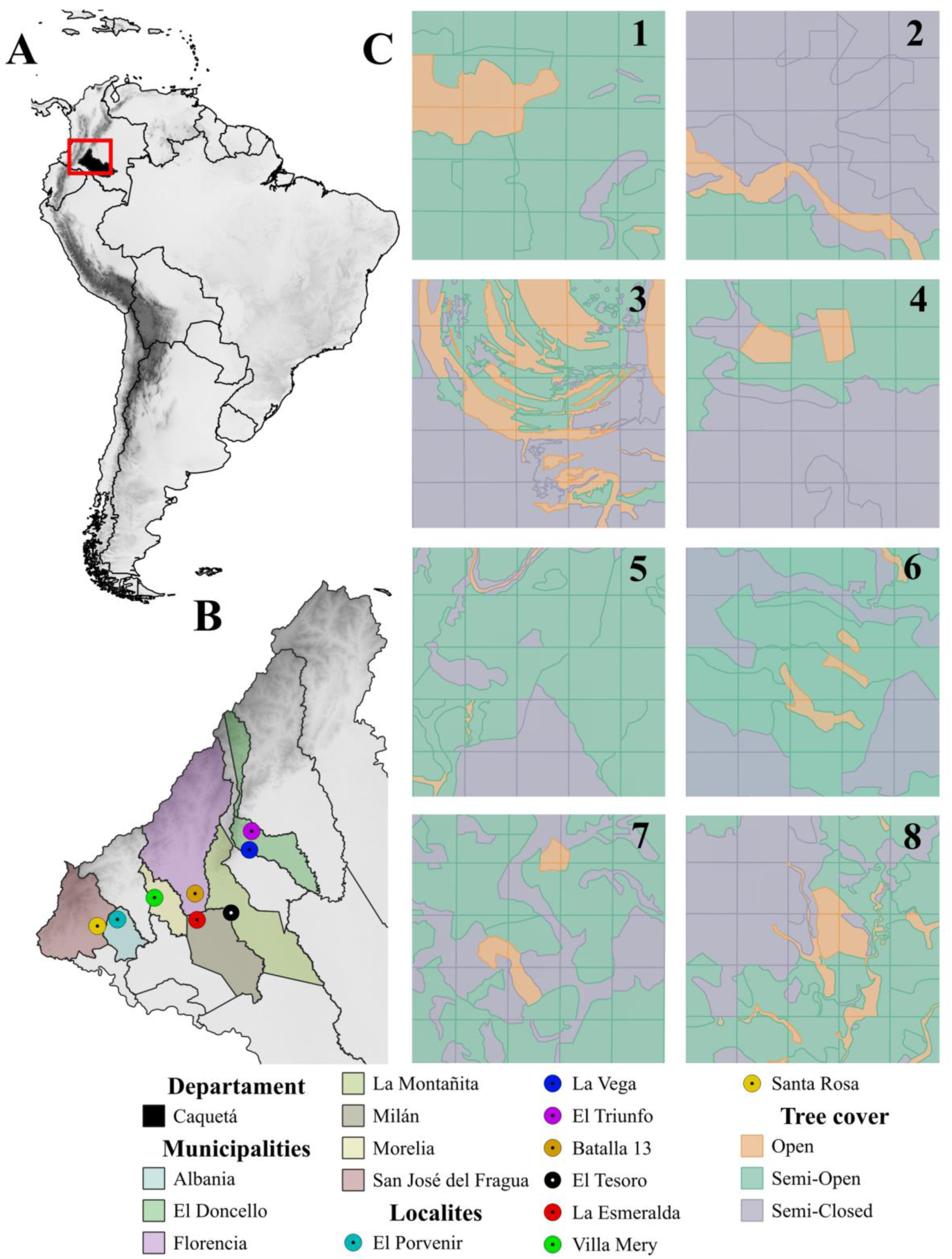
Location of the livestock landscape mosaics in the Colombian Amazon. Location of the Caquetá department at the national (Colombia) and continental (South America) scale (A); distribution of the eight mosaics across the municipalities of Caquetá (B); classification of the three types of tree cover in the studied mosaics by locality (C); Localities: El Triunfo-TR (1); La Vega-VE (2); La Esmeralda-ES (3); El Tesoro-TE (4); Villa Mery-VM (5); El Porvenir-PO (6); Santa Rosa-SR (7); Batalla 13-BA (8).

### Classification of tree cover in livestock landscape mosaics

Each landscape mosaic was defined as a square area of 1.56Lkm² (156Lha), subdivided into 25 symmetrical quadrants of 0.0625Lkm² (6.25Lha) each (Fig 2). A thematic vegetation cover map was constructed for each quadrant using the CORINE Land Cover methodology adapted for Colombia at a 1:100 000 scale [48], and for the Amazon region at a 1:25 000 scale [49]. Spatial data were obtained from geospatial vector layers downloaded from the Geoportal of the Territorial Environmental Information System of the Colombian Amazon (SIAT-AC), using 2024 data under the “Land Cover” category.

**Fig 2.**
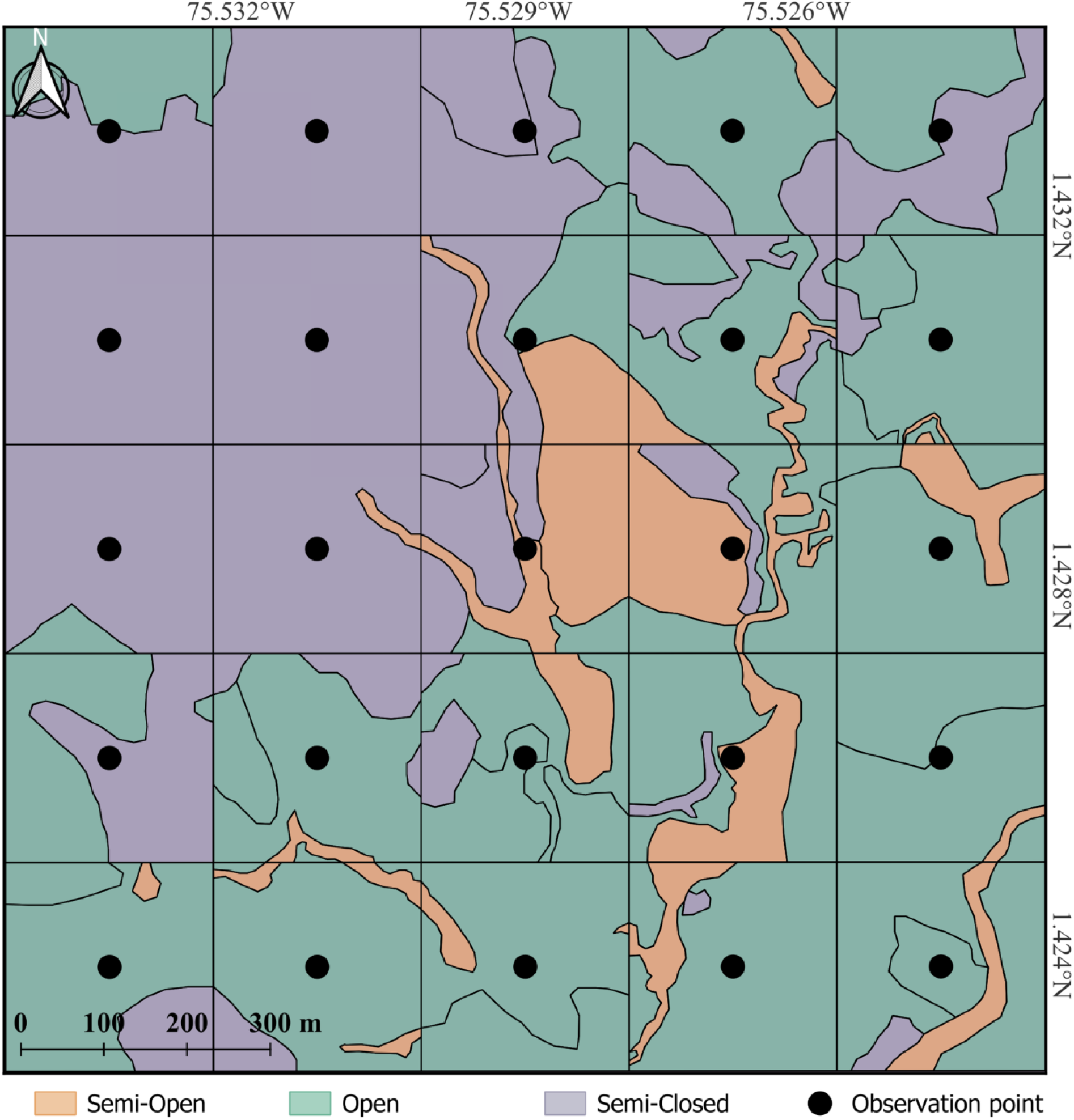
**Representation of a mosaic showing the definition of quadrants and the classification of tree cover types, along with bird census observation points in livestock landscapes in the Colombian Amazon.**

The data were validated through direct field observations and analysis of SENTINEL-2A satellite imagery, freely accessible via The Copernicus Data Space Ecosystem Services (CDSE) [50]. This sensor provides high spatial and temporal resolution multispectral images, enabling accurate detection and analysis of land cover changes in the study area [51]. Images were used at a 1:10 000 scale, with less than 10% cloud cover, ensuring high-quality spatial interpretation [52].

All images were adjusted to match the sampling units so that only data corresponding to each mosaic and quadrant within the livestock landscapes were extracted. The generated vector data were compared with reference land cover maps specific to the Amazon region to ensure the accuracy and timeliness of the information. Patch digitization and classification were carried out, and the resulting data were transformed into Shapefile format and georeferenced using the national coordinate system MAGNA-SIRGAS 2018 (EPSG:9377), via ArcMap software version 10.8 [53].

For each quadrant, the area corresponding to each type of land cover patch was calculated using QGIS software version 3.40 [54]. Subsequently, the percentage of tree cover per patch was estimated and classified into three categories according to IDEAM [48]: OP, SO, and SC (Table 1). The area of each patch was multiplied by its respective percentage of tree cover and then divided by the total area of the quadrant, allowing the total percentage of tree cover for each category within each analysis unit to be calculated. The corresponding values for each quadrant across the eight evaluated mosaics are presented in Table S1 of the supplementary material.

**Table 1.**
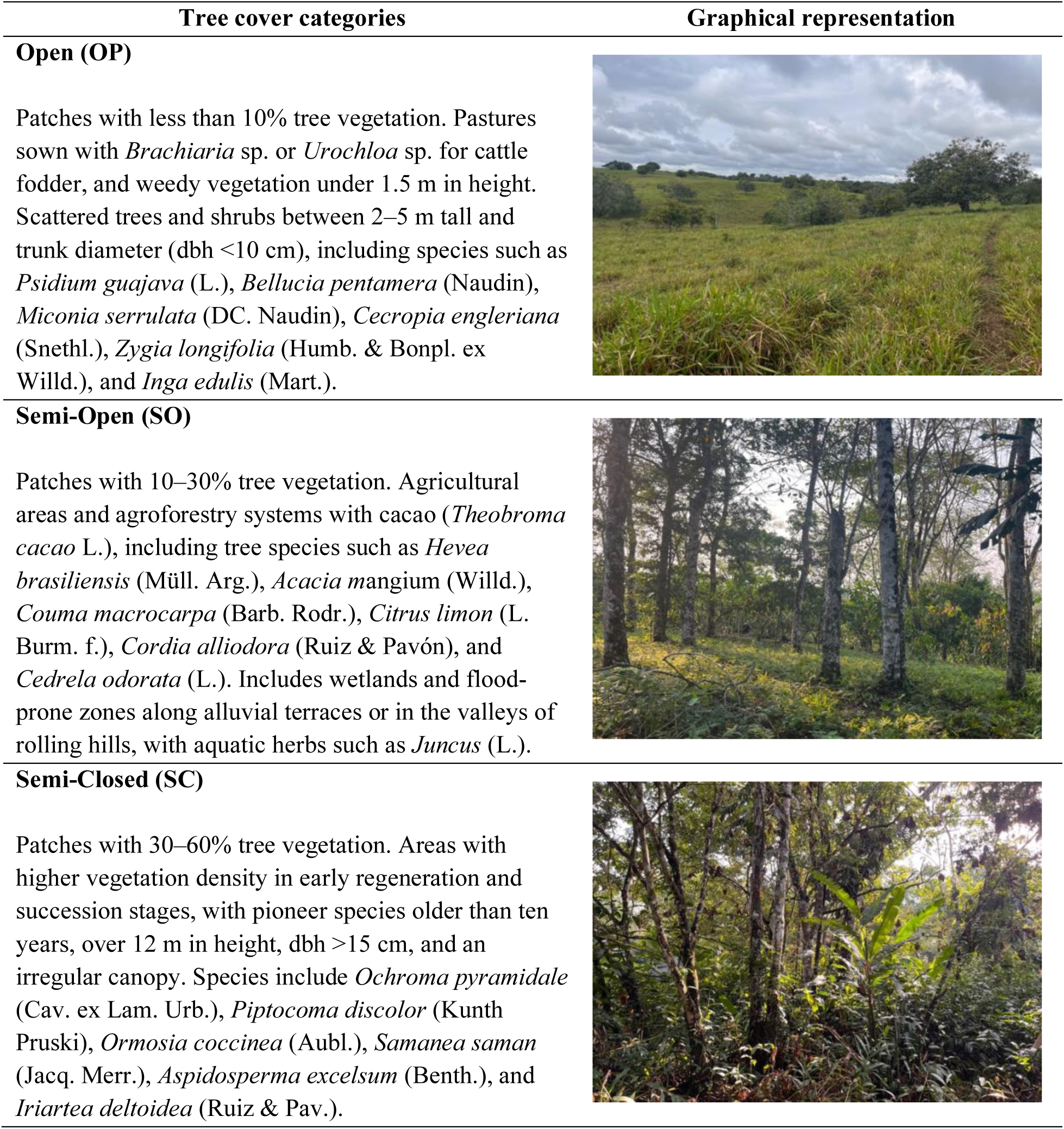
Description of tree cover categories classified within the quadrants of livestock landscape mosaics in the Colombian Amazon.

| Tree cover categories | Graphical representation |
| --- | --- |
| **Open (OP)**<br>Patches with less than 10% tree vegetation. Pastures sown with *Brachiaria* sp. or *Urochloa* sp. for cattle fodder, and weedy vegetation under 1.5 m in height. Scattered trees and shrubs between 2–5 m tall and trunk diameter (dbh <10 cm), including species such as *Psidium guajava* (L.), *Bellucia pentamera* (Naudin), *Miconia serrulata* (DC. Naudin), *Cecropia engleriana* (Snethl.), *Zygia longifolia* (Humb. & Bonpl. ex Willd.), and *Inga edulis* (Mart.). |  |
| **Semi-Open (SO)**<br>Patches with 10–30% tree vegetation. Agricultural areas and agroforestry systems with cacao (*Theobroma cacao* L.), including tree species such as *Hevea brasiliensis* (Müll. Arg.), *Acacia mangium* (Willd.), *Couma macrocarpa* (Barb. Rodr.), *Citrus limon* (L. Burm. f.), *Cordia alliodora* (Ruiz & Pavón), and *Cedrela odorata* (L.). Includes wetlands and flood-prone zones along alluvial terraces or in the valleys of rolling hills, with aquatic herbs such as *Juncus* (L.). |  |
| **Semi-Closed (SC)**<br>Patches with 30–60% tree vegetation. Areas with higher vegetation density in early regeneration and succession stages, with pioneer species older than ten years, over 12 m in height, dbh >15 cm, and an irregular canopy. Species include *Ochroma pyramidale* (Cav. ex Lam. Urb.), *Piptocoma discolor* (Kunth Pruski), *Ormosia coccinea* (Aubl.), *Samanea saman* (Jacq. Merr.), *Aspidosperma excelsum* (Benth.), and *Iriartea deltoidea* (Ruiz & Pav.). |  |

## Methods

### Bird census and functional traits

The bird diversity census was conducted between January and November 2023 using mist netting and point count methods. Observation points were established at the center of each quadrant, spaced 250 meters apart, for a total of 25 points (Fig 2). At each point, all species seen or heard were recorded over a 25-minute period during morning hours from 06:00 to 10:00, amounting to a total of 83.3 observer-hours across all landscape mosaics.

For mist netting, four 12-meter nets were set up in each quadrant, strategically positioned to intercept the flight paths of birds moving between tree-covered areas [17,55]. Nets were opened for five hours in the morning (06:00–11:00) and checked every 20 minutes to minimize stress on captured individuals. The total mist-netting effort per mosaic was 100 net-hours.

Each captured individual was measured for eight morphological traits to assess functional bird diversity (Fig 3). The traits measured included: total body length (LTO), tail length (LCO), tarsus length (LTA), extended wing length (AEX), commissure length (COM), bill height (ALT), total culmen length (CTO), and weight (PES). For birds observed but not captured, the trait matrix was completed using measurements from specimens in the Ornithological Collection of the Natural History Museum of the Universidad de la Amazonia (UAM) and from the global AVONET database of avian morphological, ecological, and geographic data [26]. Taxonomic identification to the species level followed the nomenclature of the South American Classification Committee [56], with verification using bird guides for Colombia [57–59]. Species were classified into seven guilds: frugivores (FRU), insectivores (INS), nectarivores (NEC), granivores (GRA), folivores (FOL), scavengers (CAR), and vertebrate consumers (VER). For this study, only species classified within the granivore guild (GRA) were selected for analysis.

**Fig 3.**
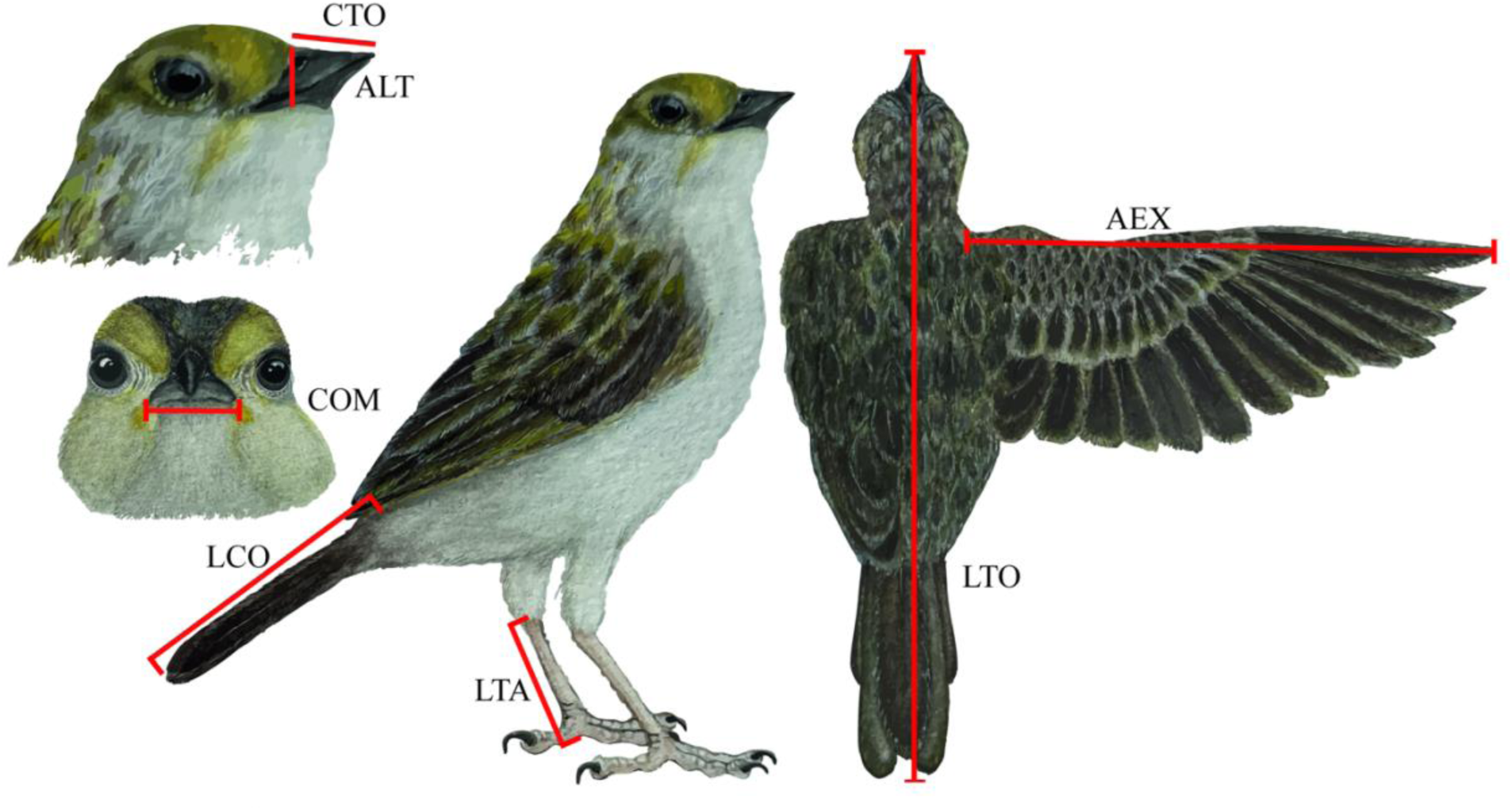
Diagram of the morphological functional traits measured per species. CTO: total culmen; LTA: tarsus length; LTO: total body length; AEX: extended wing length; LCO: tail length; COM: commissure; ALT: bill height; PES: body weight (g). Illustration of *Ammodramus aurifrons* (Passeriformes: Passerellidae) by Viviana Tello M.

### Ethics statement

The specimen collections were conducted under the framework of Resolution 1015 of 2016, in accordance with the General Permit for the Collection of Specimens of Wildlife Species for Non-Commercial Scientific Research, issued to the Universidad de la Amazonia. Approval for practices involving the use of animals was granted by the Institutional Committee on Ethics and Bioethics in Research of the Universidad de la Amazonia (FO-A-APC-01-11), established through Agreement 020 of 2018 by the Superior Council.

### Data analysis

A Generalized Linear Model (GLM) with a Poisson distribution was used to test for significant differences in the mean abundance of individuals and species richness of granivorous birds among the three categories of tree cover classified within the quadrants of livestock landscape mosaics in the Colombian Amazon. Fisher’s LSD method (α = 0.05) was applied for post-hoc comparison, using Navure statistical software, version 2.8.2 [60].

Alpha diversity was estimated using Hill numbers, which represent the effective number of species for each tree cover category. Rarefaction-extrapolation curves were plotted based on the first three Hill numbers: q0 represents species richness, q1 reflects Shannon diversity, and q2 indicates species dominance [61]. These were generated using the iNEXT.3D package in R [62].

A multivariate hierarchical cluster analysis was performed using Ward’s algorithm, with Gower’s index as the similarity measure, to group granivorous bird species based on functional trait similarity. A Principal Component Analysis (PCA) was also conducted to explore the relationship between the functional traits of granivorous bird species and the percentages of the three types of tree cover. All analyses were performed using InfoStat statistical software, version 2020 [63].

To assess differences in the functional traits of granivorous birds among the tree cover categories defined per quadrant, an analysis of variance was conducted using Linear Models (LM). The response variables were the morphometric traits of the species, and the fixed effects were the tree cover classifications: open (OP), semi-open (SO), and semi-closed (SC). Model assumptions were verified through graphical inspection of residuals. Fisher’s LSD test (α = 0.05) was applied for mean comparisons among cover types using InfoStat software, version 2020 [63].

To explore the relationship between environmental matrices (tree cover percentages) and the functional trait matrices of granivorous bird species, an RLQ analysis was conducted using the “mvabund” package [46] in R version 4.2 [64]. This multivariate approach integrates three data matrices: **R**—environmental variables, representing the percentage of open (10%), semi-open (40%), and semi-closed (60%) tree cover; **L**—species abundance matrix for granivorous birds recorded in the sampling units; and **Q**— functional trait matrix of the species. Matrices R and Q were subjected to principal component analysis (PCA), while matrix L was analyzed through correspondence analysis (COA) for abundance data. The RLQ analysis was performed using the rlq function from the ade4 package in R [65]. A co-inertia approach was applied to maximize the covariation between R and Q through L. The fourth-corner test with 9 999 permutations was used to assess the statistical significance of associations between environmental variables and functional traits, with multiple comparison correction using the False Discovery Rate (FDR) method [66].

To analyze the functional diversity of granivorous bird communities across tree cover types by quadrant, five multidimensional functional diversity indices were calculated [43,44]: FRic (functional richness), FEve (functional evenness), FDiv (functional divergence), FDis (functional dispersion), and RaoQ (quadratic entropy), using the FD package [38] in R version 4.2 [64]. An analysis of variance with Linear Models (LM) was used to test differences in the functional diversity indices among tree cover categories. The five indices (FRic, FEve, FDiv, FDis, RaoQ) were used as response variables, and the three tree cover types (OP, SO, SC) as fixed effects. Fisher’s LSD test (α = 0.05) was applied, and all analyses were carried out using InfoStat software, version 2020 [63].

To evaluate the functional diversity of granivorous bird communities across tree cover types in livestock landscapes, an AUC (Area Under the Curve) functional diversity analysis was performed [67] using Hill numbers to express the effective number of species per cover category. Rarefaction-extrapolation curves were plotted based on the first three Hill numbers (q0, q1, q2) [61,68]. This analysis was performed using the iNEXT.3D package in R [62,69].

## Results

A total of 560 individuals of granivorous birds were recorded, representing 22 species, four families, and three orders. The complete functional trait matrix of the granivorous bird species is presented in Supplementary Table S2. The order Passeriformes exhibited the highest species richness and abundance. The species *Volatinia jacarina*, *Patagioenas cayennensis*, and *Ammodramus aurifrons* accounted for approximately 50% of the individuals observed. Significant differences were found in the mean distribution of species richness (F = 3.20; p = 0.0450; df = 97) and abundance (F = 3.20; p = 0.0450; df = 97) across the tree cover types (Fig 4).

**Fig 4.**
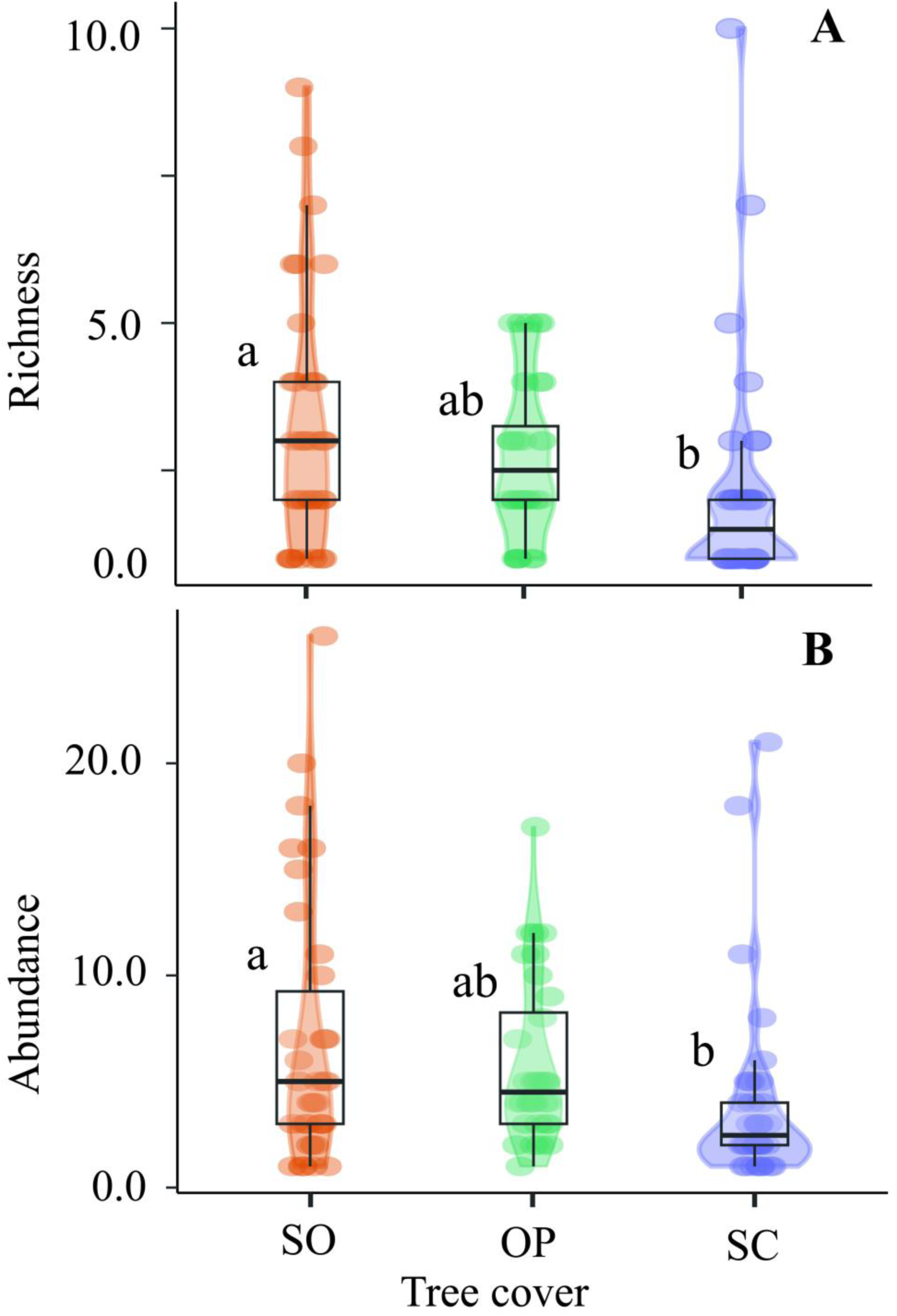
Violin plot showing the patterns of species richness (A) and abundance (B) in the granivorous bird assemblage across the tree cover gradient in livestock landscapes of the Colombian Amazon. SO: semi-open; OP: open; SC: semi-closed. Values sharing a common letter indicate that their means are not significantly different (p > 0.05).

According to Hill numbers, for index q0 (species richness), the semi-closed (SC) cover exhibited the highest number of observed species (18 spp.) from a sample of 134 individuals. Similarly, SC cover showed the highest values for Shannon diversity (q1) and Simpson diversity (q2) compared to the other cover types (Fig 5A). Sample completeness is interpreted as the number of individuals present at a site divided by the number of individuals observed at that site. In this regard, SC cover reached completeness with the fewest individuals, whereas the semi-open (SO) cover required three times as many individuals as SC to reach completeness (Fig 5B).

**Fig 5.**
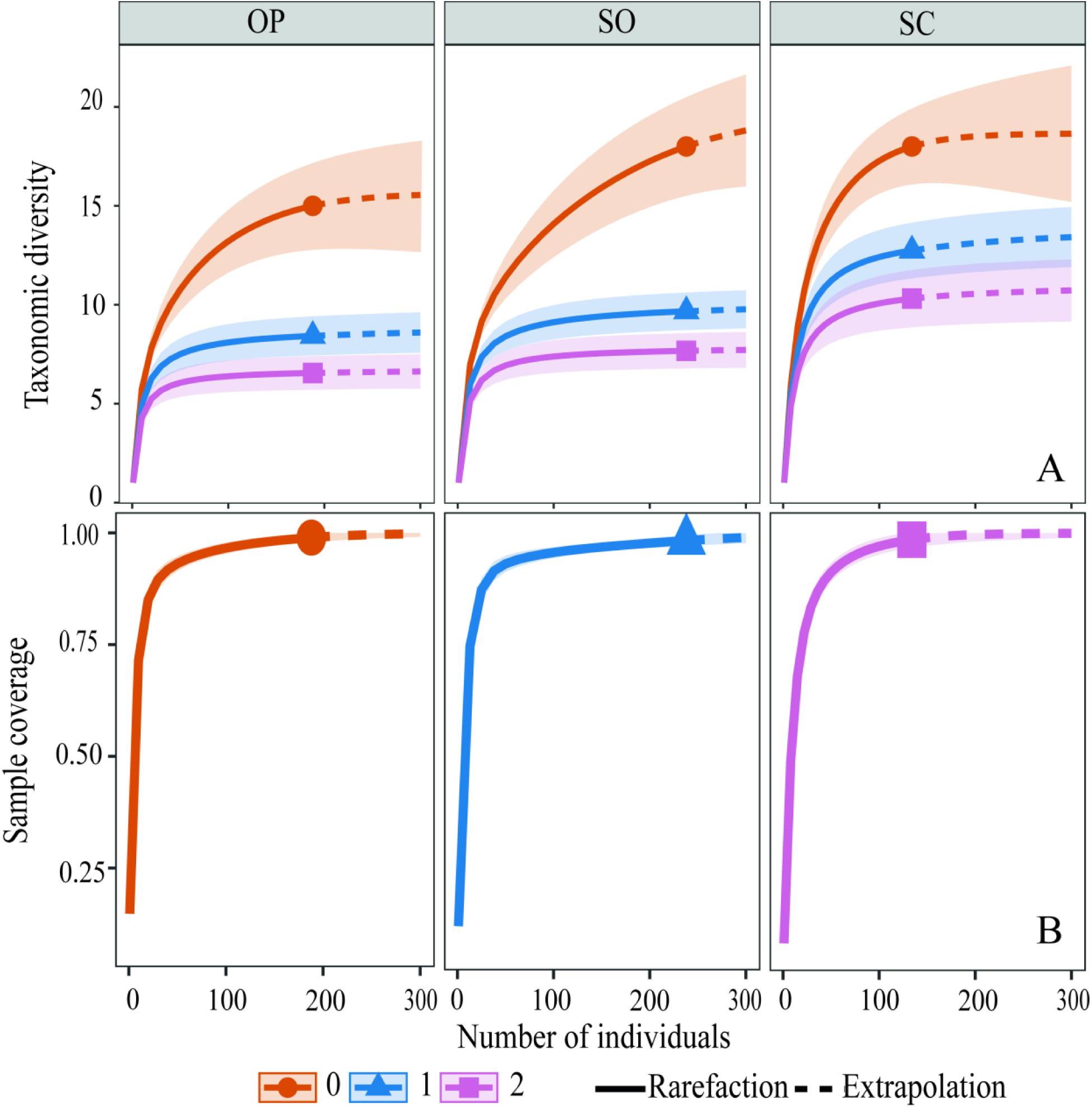
Rarefaction-extrapolation curves of diversity orders q0, q1, and q2 for the granivorous bird assemblage across the tree cover gradient in livestock landscapes of the Colombian Amazon. SO: semi-open; OP: open; SC: semi-closed. A Sample size-based sampling curve; B Sample completeness curve.

### Functional classification of species

The hierarchical cluster analysis (Gower similarity index) revealed two functional groups (Fig 6). The first group included nine granivorous species with large body sizes, belonging to the orders Columbiformes and Tinamiformes. The second group consisted of thirteen smaller species, mostly from the order Passeriformes, along with two species from the genus *Columbina*. The similarity in functional traits between these two groups was below 30%, indicating a clear separation in the morphological trait sets of large versus small granivorous birds. The species-by-group matrix resulting from the cluster analysis is presented in Supplementary Table S3.

**Fig 6.**
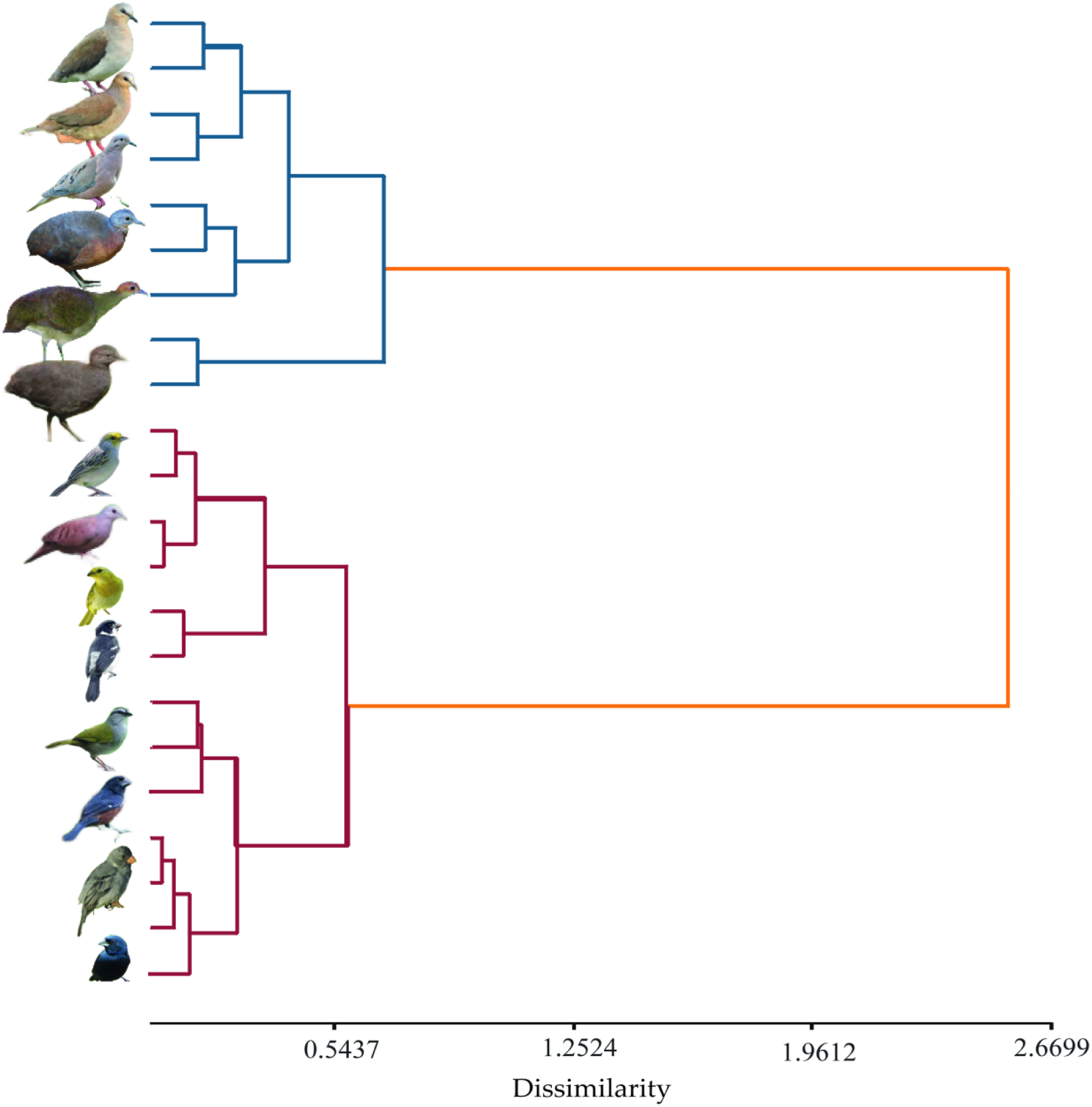
Dendrogram showing the results of the hierarchical cluster analysis using the *ward.D* algorithm. The algorithm’s distance was calculated using the default transformation of the *Gower* similarity index.

The principal component analysis (PCA) of species’ morphological traits and tree cover percentages in the quadrants of the mosaic landscapes explained 71.8% of the total variance across the first two axes. In the primary axis (PC1), the positive end grouped larger-bodied, heavier species associated with higher tree cover (semi-closed sites), while the negative end grouped smaller species with broader and taller bills, linked to open and semi-open covers (Fig 7). The second component (PC2) reflected a vegetation complexity gradient: the positive end grouped long-billed, heavy-bodied species associated with semi-open and semi-closed covers, while the negative end included species with elongated traits (long bill, extended tarsus) associated with open covers. These patterns suggest that larger-bodied species with specialized morphological traits tend to inhabit areas with denser tree cover, whereas smaller-bodied species predominate in more open environments.

**Fig 7.**
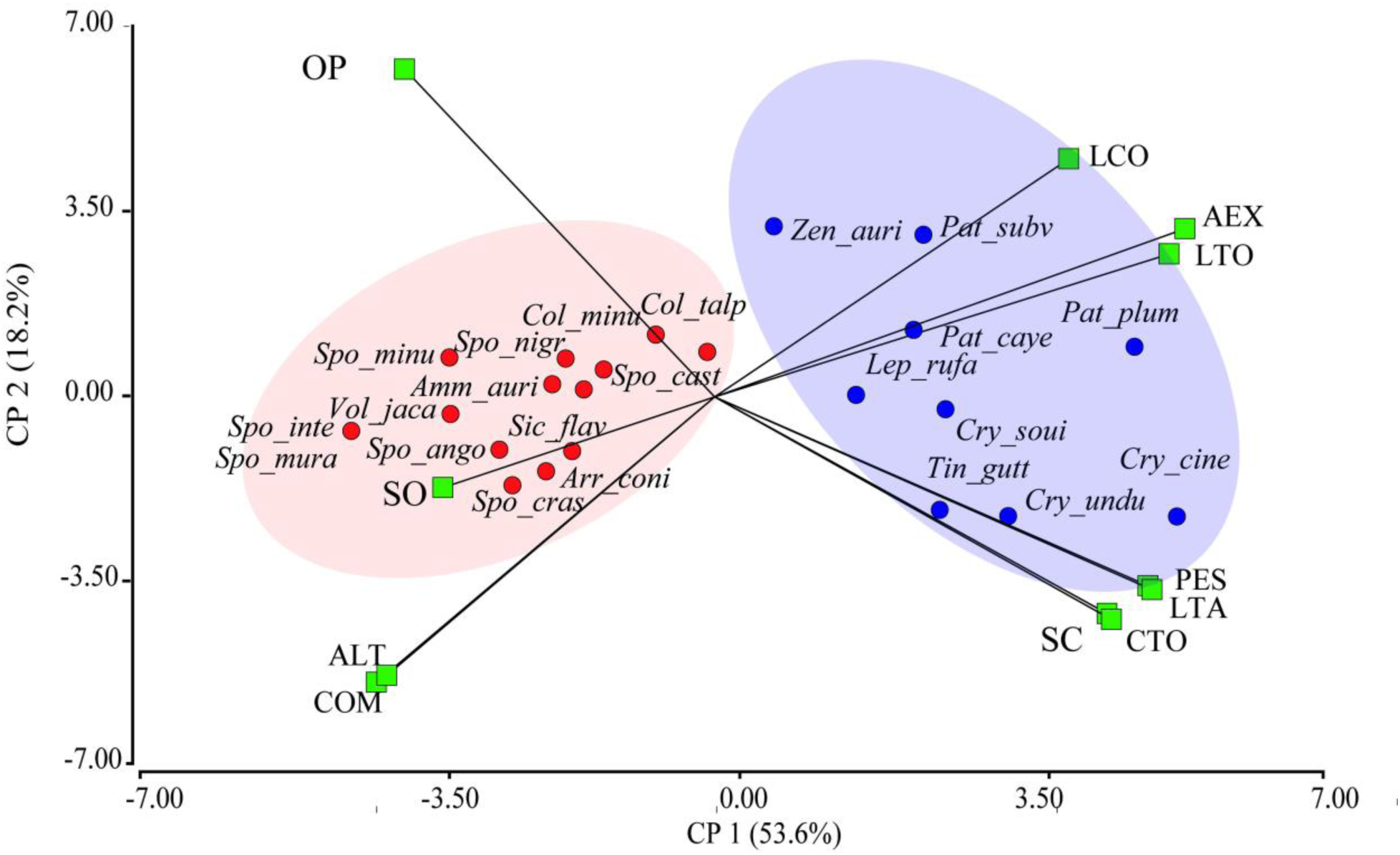
Biplot derived from the principal component analysis of species traits in the granivorous bird assemblage and tree cover types across productive landscapes in the southeastern Colombian Amazon. SO: semi-open; OP: open; SC: semi-closed; CTO: culmen total; PES: body weight (g); LTA: tarsus length; LTO: total body length; AEX: extended wing length; LCO: tail length; COM: commissure; ALT: bill height.

The functional traits of granivorous birds—culmen total (CTO; F = 6.69; p = 0.0015; n = 272; df = 269), tarsus length (LTA; F = 8.47; p = 0.0003), extended wing length (AEX; F = 3.56; p = 0.0297), total body length (LTO; F = 3.82; p = 0.0232), and body weight (PES; F = 6.52; p = 0.0017)—showed significant differences across the tree cover types, with the highest average values found in semi-closed covers.

These results indicate that tree cover influences morphological differentiation within the assemblage. The results from the Linear Model Analysis are provided in Supplementary Table S4.

### Trait–Cover association (RLQ and Fourth-Corner Analysis)

The RLQ analysis related the environmental matrix of tree cover percentage (open, semi-open, semi-closed) with the matrix of functional traits of the species (CTO, LTO, LTA, AEX, PES), weighted by species abundances. The RLQ yielded a total inertia of 0.3124, with the first axis (RLQ1) accounting for 93.42% of the projected variance and the second axis (RLQ2) explaining 6.58%. This indicates that the majority of the co-structure between functional traits and environmental variables is concentrated in the primary axis (Fig 8). The RLQ analysis reveals a functional structure in the granivorous bird communities along the gradient from open to semi-closed tree cover in cattle ranching landscapes. Species with higher values for traits such as body size, and longer bills and tarsi, were strongly associated with areas of higher tree cover. In contrast, smaller species with broader and taller bills were grouped at the opposite end of the RLQ1 axis, suggesting greater tolerance to open or disturbed environments.

**Fig 8.**
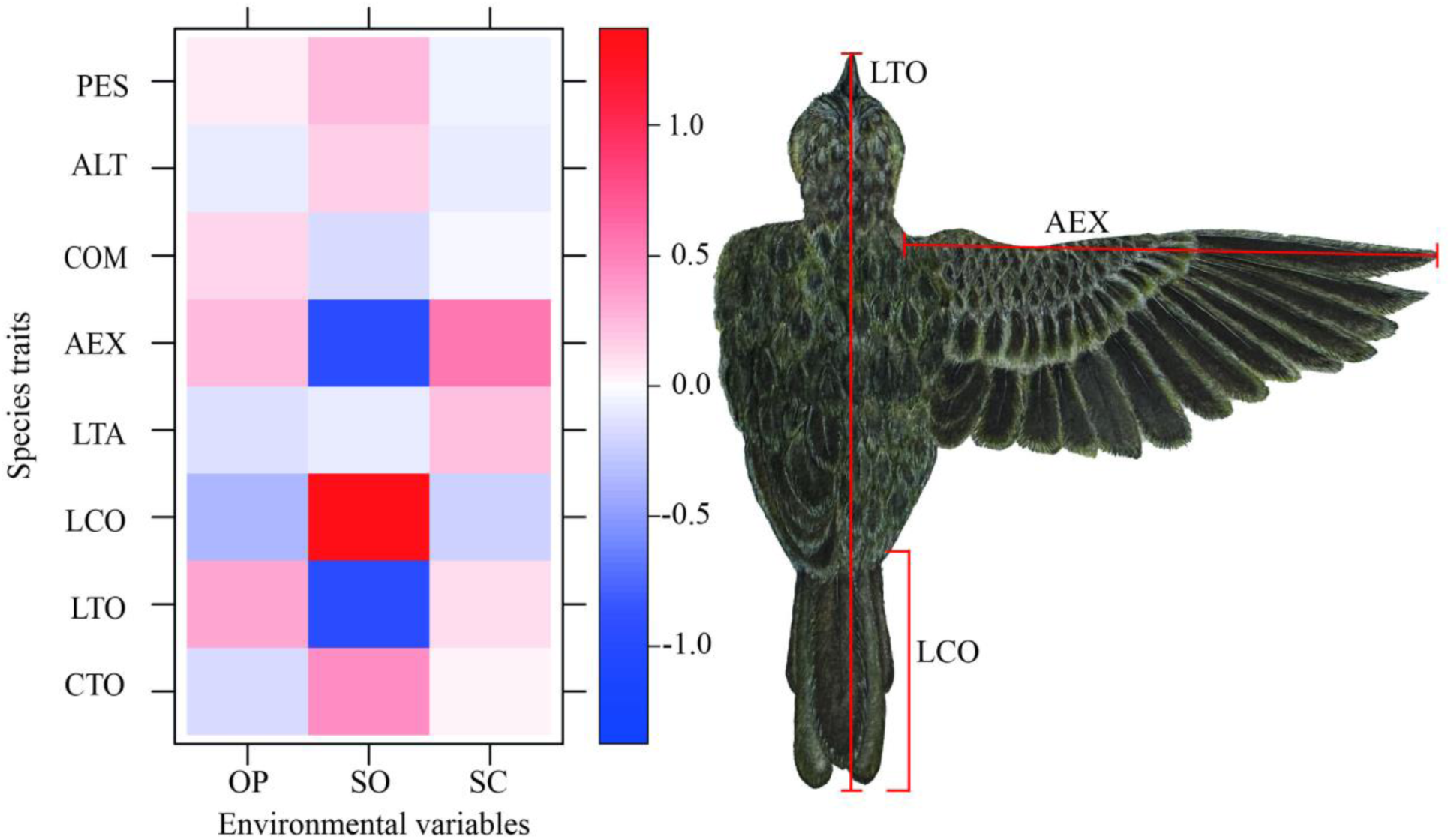
RLQ analysis between the environmental matrices of the percentage of the three types of tree cover and the functional traits of granivorous bird species in cattle ranching landscapes of the Colombian Amazon. ALT: bill height; COM: commissure; AEX: extended wing; LTA: tarsus length; LCO: tail length; LTO: total body length; CTO: total culmen; PES: body weight (g); SO: semi-open; OP: open; SC: semi-closed. Red squares indicate positive associations, while blue squares indicate negative associations. Lighter colors indicate weaker associations, whereas darker colors represent stronger associations. The illustration of *Ammodramus aurifrons* highlights the traits with the strongest associations.

These patterns were statistically confirmed by the fourth-corner test. Significant correlations (p < 0.05, FDR-adjusted) were identified between tree cover percentage and functional traits. Notably, there was a significant negative association between RLQ1 and the traits culmen total (CTO; r = –0.1756; adjusted p = 0.0496) and tarsus length (LTA; r = –0.1867; adjusted p = 0.0496), indicating that birds with longer bills and tarsi are associated with areas of denser tree cover (SC). Body weight showed a similar trend (r = –0.1723; adjusted p = 0.081, marginal), suggesting a possible relationship with more conserved habitats. Additionally, semi-closed cover (SC) was negatively correlated with RLQ1 (r = –0.1639; adjusted p = 0.0285), while open cover (OP) showed a positive correlation (r = 0.1668; adjusted p = 0.0502). Along the second axis, RLQ2, semi-open cover (SO) displayed a significant negative relationship (r = –0.1656; adjusted p = 0.0285). These associations indicate that granivorous bird species with more specialized morphological traits—such as heavier bodies and longer bills and tarsi—are linked to areas with denser tree cover, whereas open environments favor species with more generalist traits.

### Functional diversity of the granivorous bird assemblage

Regarding the mean distribution of the functional diversity indices of the granivorous bird assemblage across the quadrants of the mosaics, only the indices FDis (F = 7.12; p = 0.0013; n = 100, df = 2) and RaoQ (F = 7.41; p = 0.0010; n = 100, df = 2) exhibited significant differences among the tree cover categories, with the highest average values recorded in semi-open covers. The indices FRic, FEve, and RaoQ were highest in semi-closed cover, while FEve and FDis peaked in semi-open cover. The results of the Linear Models (LM) analysis are presented in Table S5 of the supplementary material.

In terms of AUC functional diversity, the rarefaction–extrapolation curves indicated that with a reduced number of individuals, semi-closed cover exhibited the highest values for q0, q1, and q2 (Fig 9). When extrapolated to larger sampling sizes, the functional diversity indices tended to increase, particularly in open and semi-closed covers. These findings reflect that the structural complexity of the habitat influences the levels of functional diversity within the granivorous bird assemblage.

**Fig 9.**
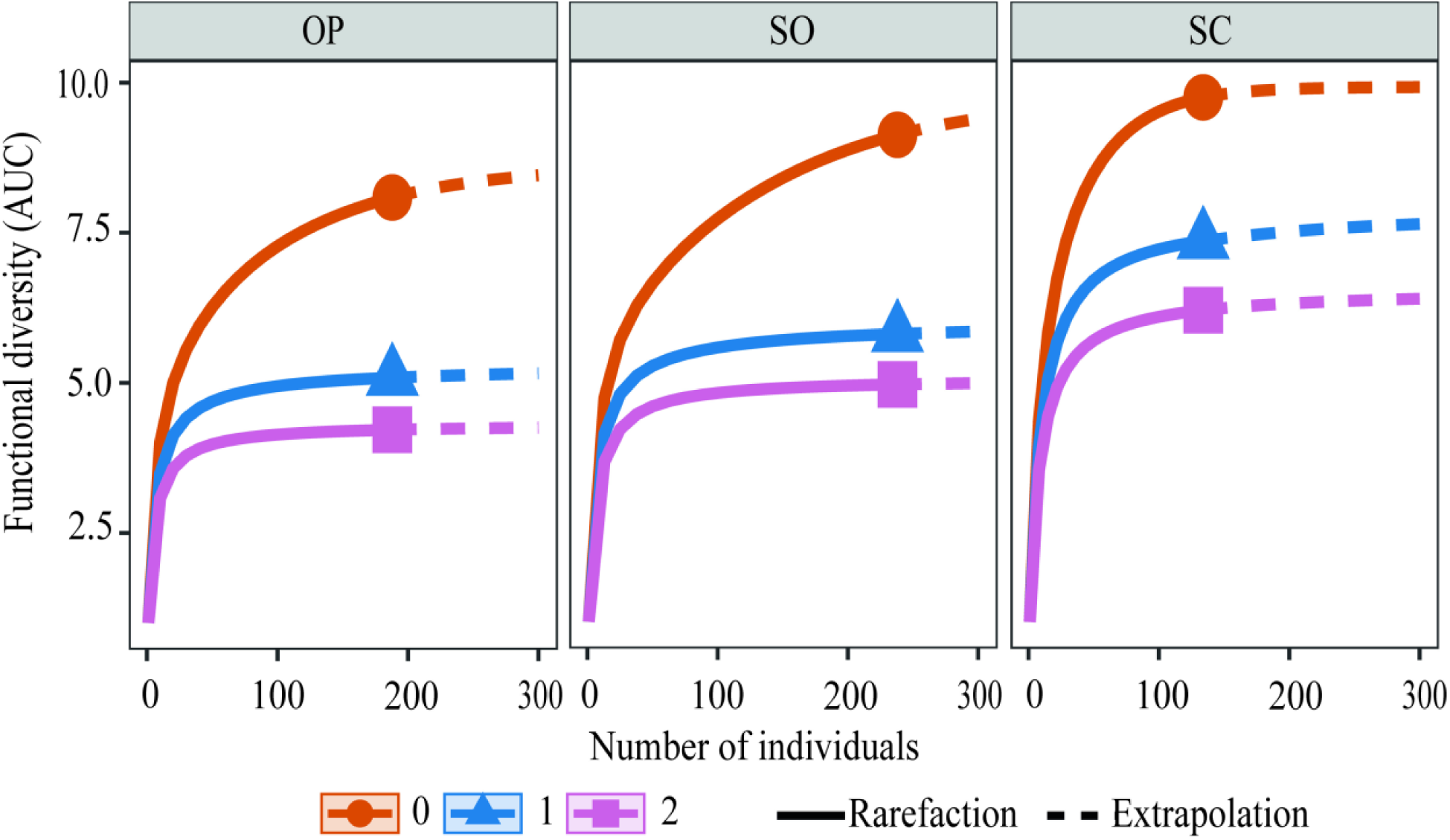
Rarefaction–extrapolation curves of functional diversity of order q0, q1, and q2 for the granivorous bird assemblage across the tree cover gradient in cattle ranching landscapes of the Colombian Amazon. SO: semi-open; OP: open; SC: semi-closed. A. Sample size-based sampling curve. B. Sample completeness curve.

## Discussion

Significant differences in species richness and abundance were observed in the granivorous bird assemblage across the different vegetation cover types. The highest richness and abundance were recorded in semi-closed and semi-open mosaics, while open-cover mosaics exhibited low taxonomic diversity and greater species homogenization. According to Echeverry et al. [70], intermediate vegetation covers tend to host higher species richness because they provide resources for both forest-dwelling and open-area species [71]. *Ammodramus aurifrons*, *Volatinia jacarina*, and *Sicalis flaveola* were the most abundant species in the study. These birds possess traits that facilitate high-density persistence in open environments dominated by pasture and early successional vegetation [72].

The gradient of tree cover percentage influenced the presence of specific granivorous bird species. Six species were exclusive to a single cover type: *Zenaida auriculata* in open areas; *Sporophila nigricollis*, *S. intermedia*, and *S. minuta* in semi-open areas. Larger granivorous birds such as *Crypturellus cinereus* and *Tinamus guttatus* were present only in semi-closed covers, being particularly affected by the expansion of pastureland in the Colombian Amazon. Notably, *T. guttatus* is currently listed as “Near Threatened” by the IUCN [73]. Granivorous species associated with denser vegetation exhibit distinctive morphological traits such as cryptic plumage, medium to large body size, robust and conical beaks, long tarsi, and short wings—adaptations linked to foraging in structurally complex habitats [74]. Such trophic and spatial specialization makes these species more sensitive to habitat fragmentation and loss [75–77].

The hierarchical clustering analysis of species’ morphological traits revealed two functional groups of granivorous birds in the cattle ranching landscapes of the Colombian Amazon. The first group consisted of larger-bodied species such as *C. cinereus*, *T. guttatus*, and *Patagioenas plumbea*. The second group included smaller-bodied species with proportionally wider and taller beaks, particularly those of the *Sporophila* and *Columbina* genera. The differentiation observed in the PCA and RLQ analyses supports this functional classification. Species with longer beaks and tarsi, greater total body length, and heavier weight were associated with higher tree cover habitats, whereas smaller-bodied birds with wider beaks were more common in open environments.

The association of total culmen length (CTO), tarsus length (LTA), and body weight (PES) with semi-closed habitats suggests that specialized functional traits confer adaptive advantages in complex habitats by enabling access to specific resources and facilitating movement through dense vegetation [34]. In contrast, open habitats favored species with generalist traits and greater mobility: smaller body size and weight, shorter tarsi, and more robust beaks—adaptations suited for diets based on *Poaceae* and *Cyperaceae* seeds, particularly forage grasses like *Brachiaria* sp., which dominate livestock pastures and benefit granivores adapted to open landscapes [78,79]. These findings align with previous research indicating that changes in tree density in cattle ranching landscapes lead to morphological differentiation among species and declines in avian body condition [35,80].

Overall, this study indicates that tree cover reduction in cattle ranching landscapes acts as an environmental filter that excludes species with specialized functional traits, promoting communities with lower functional diversity. Environmental degradation due to land-use changes leads to biotic homogenization at multiple spatial scales, threatening ecosystem integrity and biodiversity conservation [31]. However, biological community responses to habitat disturbance vary among species and functional guilds, depending on their resource use strategies and ecological tolerance levels [16,18,81].

Granivorous birds were distributed along the tree cover gradient, with the highest richness and abundance occurring in semi-closed and semi-open covers. According to Díaz-Cháux et al. [17], in productive Amazonian landscapes, avian functional diversity varies according to the structure and composition of vegetation in agroforestry and silvopastoral systems. In contrast, open covers favored generalist species adapted to disturbed environments. This pattern is consistent with previous studies showing that variation in tree density and landscape structural heterogeneity promotes bird diversity, spatial segregation, and interspecific coexistence [30,82–84].

### Functional diversity of the granivorous bird assemblage

Functional diversity followed a similar pattern to taxonomic diversity, with higher values of FRic, FEve, and RaoQ indices observed in granivorous bird assemblages from semi-closed covers, and higher FDiv and FDis values in semi-open covers. This pattern reveals low overlap in trophic niches across communities along the tree cover gradient, resulting from the even distribution of functionally non-redundant traits and divergent trophic strategies [22,85]. Such a condition reflects the increased structural heterogeneity of denser vegetation covers, which facilitates interspecific coexistence by increasing the availability of microhabitats and seeds of diverse morphologies [21,23]. These environments are often associated with larger-bodied species that exhibit specialized morphological traits such as longer tarsi and culmen, greater total body length, and heavier body mass.

Amazonian cattle-ranching landscapes are spatially structured by open matrices interspersed with scattered patches of remnant vegetation, characterized by low heterogeneity and limited connectivity [9,47]. In such landscapes, structural simplification reduces functional trait diversity and promotes communities that are functionally redundant and homogeneous [5,83,86]. In this regard, the present study found that open-cover communities exhibited the lowest functional diversity values, forming assemblages with high redundancy and low complementarity, which restricts the breadth of functional niches. These findings support the hypothesis that tree cover and resource availability act as environmental filters shaping both the taxonomic and functional structure of granivorous bird assemblages. The loss of functionality in highly disturbed areas serves as a warning signal about the ecological consequences of deforestation on ecosystem resilience [87–91]. This pattern is consistent with studies that document the loss of specialized traits in deforested landscapes, which may compromise key ecological functions and reduce the provision of ecosystem services such as seed dispersal, weed control, and natural vegetation regeneration [32,92–95].

Further research should integrate other avian trophic guilds using multiscale and temporal approaches that account for patterns of tree cover use and interspecific dynamics in Amazonian landscapes. Such studies would enhance our understanding of the ecological processes structuring bird communities in anthropogenically transformed environments—critical for designing management strategies that promote both ecological and economic sustainability in the region. These results underscore the need to incorporate ecological restoration strategies and sustainable land management into productive landscapes by conserving forest patches and dense vegetation corridors within cattle ranching matrices.

### Implications for Ecosystem Service provision

According to Filgueiras et al. [96] and Brocardo et al. [97], the provision of ecosystem services depends not only on species richness but also on functional complementarity and coexistence patterns at multiple spatial scales. Differentiation in functional traits among vegetation cover types enhances niche partitioning and reduces interspecific competition, allowing for the exploitation of a broader range of resources. In this context, and as proposed by Díaz-Cháux et al. [17], the granivorous bird assemblage constitutes a functional guild, given its role in similar ecological functions and its high trait variability, which translates into low functional redundancy and a significant contribution to the regulation of ecosystem services along the gradient of tree cover.

As key components of functional biodiversity in cattle ranching landscapes, granivorous birds play essential roles in seed dispersal and germination, gene flow, nutrient cycling, and the control of weeds or herbaceous invaders [98,99]. However, these species are vulnerable to the intensification of pasture expansion and the loss of tree cover [19], which threaten ecological functioning in these ranching landscapes [100–104], particularly in ecologically vulnerable regions like the Colombian Amazon.

In this context, the findings of this study serve as a warning about the ecological consequences of tree cover loss on the conservation of avian functional diversity. It is recommended to promote silvopastoral systems with mosaic configurations that include semi-open covers and dense vegetative corridors, capable of supporting species with high ecological requirements and preserving ecosystem functionality. The trait-based approach represents a robust tool for assessing biological responses to land-use change and for guiding conservation strategies focused on the multifunctionality of rural landscapes.

## Conclusions

Tree cover in cattle-ranching mosaic landscapes of the Colombian Amazon acts as an ecological filter that shapes both the taxonomic and functional diversity of granivorous bird assemblages. Habitat loss and structural homogenization reduce functional diversity, favoring communities dominated by generalist species that are functionally redundant and less capable of supporting key ecological processes. In contrast, semi-closed and semi-open vegetation covers sustain a wider variety of functional traits, promote niche partitioning, and support the coexistence of species with diverse ecological strategies. This translates into greater ecosystem resilience, enhanced multifunctionality, and increased provision of ecosystem services with both direct and indirect benefits for rural communities.

These findings underscore the importance of conserving and restoring structural heterogeneity as part of sustainable landscape management, particularly through the implementation of silvopastoral systems and the protection of native vegetation patches. Integrating a functional trait-based approach into land-use planning provides a robust framework for assessing the impact of deforestation on biodiversity and ecosystem services. It also enables the prioritization of management actions that ensure ecological and productive sustainability in environmentally vulnerable regions such as the Colombian Amazon.

Accordingly, this study reaffirms the need to promote landscape management strategies centered on the conservation of ecological traits and functions, complementing traditional biodiversity indicators such as species richness and abundance. Maintaining the functional diversity of guilds such as granivorous birds is essential for preserving core ecological processes, securing the provision of ecosystem services, and safeguarding the ecological integrity of tropical livestock production systems.

## Supporting information

Material complementario

## Acknowledgments

The authors thank the Universidad de la Amazonia (Caquetá, Colombia) and the Centro de Investigación de la Biodiversidad Andino Amazónica -INBIANAM for providing the equipment used for data collection. We also express our gratitude to the Comité Departamental de Ganaderos del Caquetá for facilitating access to the study sites, to biologist and illustrator Viviana Tello M. for the bird illustrations, and to the Semillero de Investigación en Biodiversidad y Servicios Ecosistémicos -BySE for their support during field data collection.

## Supporting information

The complete information is provided in the supplementary material file (.docx).

**Table S1**. Total percentage of tree cover for each of the three classified categories within the quadrants established across the eight livestock landscapes mosaics in the Colombian Amazon.

**Table S2**. List and matrix of morphological functional traits of the 22 granivorous bird species recorded across the three types of tree cover in livestock landscapes of the Colombian Amazon.

**Table S3**. Matrix of granivorous bird species assigned to two groups (small and large) based on the dendrogram generated by hierarchical cluster analysis using the ward.D algorithm.

**Table S4**. Linear Models Analysis and pairwise Fisher’s mean comparisons (α = 0.05) of the functional traits of granivorous bird species across three types of tree cover in livestock landscapes of the Colombian Amazon.

**Table S5**. Linear Models Analysis and pairwise Fisher’s mean comparisons (α = 0.05) of functional diversity indices of granivorous birds across three types of tree cover in livestock landscapes of the Colombian Amazon.

