## Supplementary material for "Livestock Landscapes as Ecological Filters: Effects of the Tree Cover Gradient on the Taxonomic and Functional Diversity of Granivorous Birds in the Colombian Amazon": Material complementario

**Table S1. Total percentage of tree cover for each of the three classified categories within the quadrants established across the eight livestock landscapes mosaics in the Colombian Amazon.**

| Municipalities | Mosaics | Mosaics code | Quadrant code | Tree cover | | |
|---|---|---|---|---|---|---|
| | | | | OP | SO | SC |
| Albania | El Porvenir | PO | PO03 | 0.081 | 0.000 | 0.114 |
| | | | PO04 | 0.041 | 0.000 | 0.356 |
| | | | PO05 | 0.046 | 0.000 | 0.327 |
| | | | PO06 | 0.000 | 0.000 | 0.600 |
| | | | PO07 | 0.057 | 0.030 | 0.208 |
| | | | PO09 | 0.071 | 0.003 | 0.170 |
| | | | PO11 | 0.070 | 0.000 | 0.178 |
| | | | PO12 | 0.082 | 0.022 | 0.072 |
| | | | PO13 | 0.087 | 0.047 | 0.000 |
| | | | PO14 | 0.085 | 0.036 | 0.029 |
| | | | PO15 | 0.070 | 0.000 | 0.182 |
| | | | PO16 | 0.052 | 0.000 | 0.291 |
| | | | PO17 | 0.090 | 0.035 | 0.000 |
| | | | PO18 | 0.095 | 0.006 | 0.017 |
| | | | PO19 | 0.091 | 0.000 | 0.055 |
| | | | PO22 | 0.041 | 0.000 | 0.351 |
| | | | PO23 | 0.073 | 0.000 | 0.161 |
| | | | PO24 | 0.061 | 0.019 | 0.200 |
| | | | PO25 | 0.083 | 0.025 | 0.059 |
| El Doncello | El Tesoro | TE | TE01 | 0.000 | 0.000 | 0.600 |
| | | | TE02 | 0.000 | 0.000 | 0.600 |
| | | | TE04 | 0.000 | 0.000 | 0.600 |
| | | | TE05 | 0.000 | 0.000 | 0.600 |
| | | | TE06 | 0.000 | 0.000 | 0.600 |
| | | | TE08 | 0.000 | 0.000 | 0.600 |
| | | | TE10 | 0.000 | 0.000 | 0.600 |
| | | | TE12 | 0.012 | 0.000 | 0.526 |
| | | | TE14 | 0.037 | 0.000 | 0.378 |
| | | | TE15 | 0.037 | 0.000 | 0.378 |
| | | | TE17 | 0.039 | 0.187 | 0.048 |
| | | | TE21 | 0.100 | 0.000 | 0.000 |
| | | | TE23 | 0.075 | 0.058 | 0.050 |
| | | | TE24 | 0.085 | 0.003 | 0.086 |
| | La Vega | VE | VE01 | 0.100 | 0.000 | 0.000 |
| | | | VE02 | 0.100 | 0.000 | 0.000 |
| | | | VE03 | 0.071 | 0.003 | 0.171 |
| | | | VE14 | 0.000 | 0.000 | 0.600 |
| | | | VE15 | 0.000 | 0.000 | 0.600 |
| | | | VE19 | 0.000 | 0.000 | 0.600 |
| | | | VE23 | 0.000 | 0.000 | 0.600 |
| Florencia | Batalla 13 | BA | BA01 | 0.082 | 0.006 | 0.098 |
| | | | BA03 | 0.094 | 0.022 | 0.000 |
| | | | BA04 | 0.087 | 0.042 | 0.007 |

| | | | | | | |
|---|---|---|---|---|---|---|
| | | | BA05 | 0.083 | 0.045 | 0.026 |
| | | | BA07 | 0.066 | 0.106 | 0.024 |
| | | | BA08 | 0.074 | 0.050 | 0.068 |
| | | | BA09 | 0.081 | 0.014 | 0.090 |
| | | | BA10 | 0.050 | 0.000 | 0.298 |
| | | | BA12 | 0.000 | 0.008 | 0.587 |
| | | | BA13 | 0.010 | 0.169 | 0.247 |
| | | | BA14 | 0.044 | 0.155 | 0.072 |
| | | | BA15 | 0.084 | 0.055 | 0.000 |
| | | | BA16 | 0.093 | 0.004 | 0.032 |
| | | | BA17 | 0.063 | 0.045 | 0.144 |
| | | | BA18 | 0.025 | 0.096 | 0.286 |
| | | | BA19 | 0.000 | 0.000 | 0.600 |
| La Montañita | El Triunfo | TR | TR01 | 0.100 | 0.000 | 0.000 |
| | | | TR02 | 0.100 | 0.000 | 0.000 |
| | | | TR03 | 0.100 | 0.000 | 0.000 |
| | | | TR04 | 0.086 | 0.000 | 0.085 |
| | | | TR06 | 0.088 | 0.000 | 0.075 |
| | | | TR07 | 0.066 | 0.000 | 0.203 |
| | | | TR08 | 0.100 | 0.000 | 0.000 |
| | | | TR09 | 0.100 | 0.000 | 0.000 |
| | | | TR10 | 0.100 | 0.000 | 0.000 |
| | | | TR11 | 0.045 | 0.192 | 0.000 |
| | | | TR13 | 0.093 | 0.026 | 0.000 |
| | | | TR14 | 0.100 | 0.000 | 0.001 |
| | | | TR15 | 0.091 | 0.000 | 0.052 |
| | | | TR16 | 0.092 | 0.000 | 0.051 |
| | | | TR17 | 0.093 | 0.000 | 0.042 |
| | | | TR18 | 0.067 | 0.115 | 0.000 |
| | | | TR19 | 0.026 | 0.258 | 0.000 |
| | | | TR20 | 0.032 | 0.236 | 0.000 |
| | | | TR21 | 0.100 | 0.000 | 0.000 |
| | | | TR22 | 0.100 | 0.000 | 0.000 |
| | | | TR24 | 0.100 | 0.000 | 0.000 |
| | | | TR25 | 0.100 | 0.000 | 0.000 |
| Milán | La Esmeralda | ES | ES02 | 0.000 | 0.000 | 0.600 |
| | | | ES03 | 0.000 | 0.034 | 0.542 |
| | | | ES06 | 0.000 | 0.027 | 0.553 |
| | | | ES07 | 0.000 | 0.109 | 0.412 |
| | | | ES12 | 0.028 | 0.212 | 0.067 |
| | | | ES14 | 0.032 | 0.090 | 0.254 |
| Morelia | Villa Mery | VM | VM01 | 0.080 | 0.000 | 0.119 |
| | | | VM02 | 0.088 | 0.000 | 0.074 |
| | | | VM03 | 0.070 | 0.000 | 0.178 |
| | | | VM04 | 0.098 | 0.000 | 0.012 |
| | | | VM05 | 0.072 | 0.037 | 0.103 |
| | | | VM07 | 0.073 | 0.023 | 0.123 |
| | | | VM08 | 0.067 | 0.000 | 0.200 |

| | | | | | | |
|---|---|---|---|---|---|---|
| | | | VM17 | 0.089 | 0.000 | 0.067 |
| | | | VM18 | 0.100 | 0.000 | 0.000 |
| | | | VM19 | 0.100 | 0.000 | 0.000 |
| San José del Fragua | Santa Rosa | SR | SR02 | 0.078 | 0.000 | 0.132 |
| | | | SR05 | 0.046 | 0.000 | 0.325 |
| | | | SR06 | 0.082 | 0.000 | 0.109 |
| | | | SR15 | 0.034 | 0.000 | 0.394 |
| | | | SR18 | 0.025 | 0.076 | 0.320 |
| | | | SR21 | 0.019 | 0.000 | 0.486 |

OP: open; SO: semi-open; SC: semi-closed.

**Table S2. List and matrix of morphological functional traits of the 22 granivorous bird species recorded across the three types of tree cover in livestock landscapes of the Colombian Amazon.**

| Order | Family | Scientific name | Mosaics | Quadrant | TC | AB | Morphological functional traits | | | | | | | |
|---|---|---|---|---|---|---|---|---|---|---|---|---|---|---|
| | | | | | | | CTO | LTO | LCO | LTA | AEX | COM | ALT | PES |
| Columbiformes | Columbidae | *Columbina minuta* | BA | BA07 | SO | 1 | 15.25 | 147.00 | 57.60 | 12.50 | 119.00 | 10.91 | 3.71 | 31.10 |
| | | | BA | BA16 | OP | 4 | 16.03 | 148.08 | 55.22 | 11.75 | 122.63 | 5.99 | 3.43 | 35.47 |
| | | | TE | TE05 | SC | 4 | 16.03 | 148.08 | 55.22 | 11.75 | 122.63 | 5.99 | 3.43 | 35.47 |
| | | | TE | TE15 | SC | 3 | 16.03 | 148.08 | 55.22 | 11.75 | 122.63 | 5.99 | 3.43 | 35.47 |
| | | | TR | TR03 | OP | 1 | 16.03 | 148.08 | 55.22 | 11.75 | 122.63 | 5.99 | 3.43 | 35.47 |
| | | | TR | TR15 | OP | 1 | 20.17 | 141.03 | 54.39 | 10.17 | 114.83 | 4.05 | 3.70 | 34.20 |
| | | | VM | VM07 | SO | 1 | 12.66 | 156.20 | 53.67 | 12.59 | 134.06 | 3.02 | 2.87 | 41.10 |
| | | *Columbina talpacoti* | BA | BA01 | SO | 1 | 18.01 | 158.01 | 65.02 | 12.79 | 133.89 | 4.54 | 4.43 | 43.18 |
| | | | BA | BA12 | SC | 4 | 18.01 | 158.01 | 65.02 | 12.79 | 133.89 | 4.54 | 4.43 | 43.18 |
| | | | BA | BA14 | SO | 4 | 18.57 | 163.01 | 68.39 | 12.78 | 131.95 | 4.35 | 4.16 | 40.49 |
| | | | BA | BA16 | OP | 5 | 18.01 | 158.01 | 65.02 | 12.79 | 133.89 | 4.54 | 4.43 | 43.18 |
| | | | PO | PO11 | SO | 4 | 18.01 | 158.01 | 65.02 | 12.79 | 133.89 | 4.54 | 4.43 | 43.18 |
| | | | PO | PO23 | SO | 5 | 18.01 | 158.01 | 65.02 | 12.79 | 133.89 | 4.54 | 4.43 | 43.18 |
| | | | PO | PO24 | SO | 1 | 18.01 | 158.01 | 65.02 | 12.79 | 133.89 | 4.54 | 4.43 | 43.18 |
| | | | TE | TE05 | SC | 2 | 18.01 | 158.01 | 65.02 | 12.79 | 133.89 | 4.54 | 4.43 | 43.18 |
| | | | TE | TE10 | SC | 1 | 18.01 | 158.01 | 65.02 | 12.79 | 133.89 | 4.54 | 4.43 | 43.18 |
| | | | TE | TE15 | SC | 1 | 18.01 | 158.01 | 65.02 | 12.79 | 133.89 | 4.54 | 4.43 | 43.18 |
| | | | TE | TE17 | SO | 2 | 18.01 | 158.01 | 65.02 | 12.79 | 133.89 | 4.54 | 4.43 | 43.18 |
| | | | TE | TE21 | OP | 2 | 18.01 | 158.01 | 65.02 | 12.79 | 133.89 | 4.54 | 4.43 | 43.18 |
| | | | TE | TE23 | SO | 3 | 18.01 | 158.01 | 65.02 | 12.79 | 133.89 | 4.54 | 4.43 | 43.18 |
| | | | TE | TE24 | SO | 4 | 18.01 | 158.01 | 65.02 | 12.79 | 133.89 | 4.54 | 4.43 | 43.18 |
| | | | TR | TR13 | OP | 4 | 18.01 | 158.01 | 65.02 | 12.79 | 133.89 | 4.54 | 4.43 | 43.18 |
| | | | TR | TR18 | SO | 1 | 20.66 | 142.04 | 64.18 | 9.07 | 132.48 | 4.53 | 4.46 | 38.70 |
| | | | TR | TR20 | SO | 3 | 18.01 | 158.01 | 65.02 | 12.79 | 133.89 | 4.54 | 4.43 | 43.18 |
| | | | VE | VE19 | SC | 2 | 16.13 | 161.00 | 62.07 | 14.66 | 136.53 | 4.74 | 4.68 | 48.10 |
| | | *Leptotila rufaxilla* | BA | BA12 | SC | 2 | 27.37 | 240.33 | 89.20 | 22.76 | 187.33 | 6.21 | 5.08 | 119.27 |
| | | | BA | BA16 | OP | 2 | 27.37 | 240.33 | 89.20 | 22.76 | 187.33 | 6.21 | 5.08 | 119.27 |
| | | | ES | ES06 | SC | 1 | 27.60 | 240.00 | 88.90 | 22.30 | 187.00 | 5.98 | 5.01 | 118.90 |
| | | | ES | ES07 | SC | 1 | 26.50 | 235.00 | 86.70 | 21.98 | 185.00 | 5.76 | 4.99 | 119.20 |

| | | | | | | | | | | | | |
|---|---|---|---|---|---|---|---|---|---|---|---|---|
| | PO | PO09 | SO | 2 | 27.37 | 240.33 | 89.20 | 22.76 | 187.33 | 6.21 | 5.08 | 119.27 |
| | PO | PO11 | SO | 2 | 27.37 | 240.33 | 89.20 | 22.76 | 187.33 | 6.21 | 5.08 | 119.27 |
| | PO | PO23 | SO | 2 | 27.37 | 240.33 | 89.20 | 22.76 | 187.33 | 6.21 | 5.08 | 119.27 |
| | TR | TR11 | SO | 1 | 28.00 | 246.00 | 92.00 | 24.00 | 190.00 | 6.88 | 5.23 | 119.70 |
| | TR | TR16 | OP | 1 | 27.37 | 240.33 | 89.20 | 22.76 | 187.33 | 6.21 | 5.08 | 119.27 |
| *Patagioenas cayennensis* | BA | BA01 | SO | 2 | 24.50 | 245.00 | 112.70 | 23.40 | 258.00 | 4.60 | 5.50 | 229.00 |
| | BA | BA03 | OP | 12 | 24.50 | 245.00 | 112.70 | 23.40 | 258.00 | 4.60 | 5.50 | 229.00 |
| | BA | BA07 | SO | 2 | 24.50 | 245.00 | 112.70 | 23.40 | 258.00 | 4.60 | 5.50 | 229.00 |
| | BA | BA08 | SO | 2 | 24.50 | 245.00 | 112.70 | 23.40 | 258.00 | 4.60 | 5.50 | 229.00 |
| | BA | BA09 | SO | 1 | 24.50 | 245.00 | 112.70 | 23.40 | 258.00 | 4.60 | 5.50 | 229.00 |
| | BA | BA14 | SO | 2 | 24.50 | 245.00 | 112.70 | 23.40 | 258.00 | 4.60 | 5.50 | 229.00 |
| | BA | BA15 | OP | 1 | 24.50 | 245.00 | 112.70 | 23.40 | 258.00 | 4.60 | 5.50 | 229.00 |
| | BA | BA17 | SO | 2 | 24.50 | 245.00 | 112.70 | 23.40 | 258.00 | 4.60 | 5.50 | 229.00 |
| | BA | BA18 | SC | 4 | 24.50 | 245.00 | 112.70 | 23.40 | 258.00 | 4.60 | 5.50 | 229.00 |
| | BA | BA19 | SC | 2 | 24.50 | 245.00 | 112.70 | 23.40 | 258.00 | 4.60 | 5.50 | 229.00 |
| | PO | PO03 | SO | 2 | 24.50 | 245.00 | 112.70 | 23.40 | 258.00 | 4.60 | 5.50 | 229.00 |
| | PO | PO04 | SC | 1 | 24.50 | 245.00 | 112.70 | 23.40 | 258.00 | 4.60 | 5.50 | 229.00 |
| | PO | PO05 | SC | 2 | 24.50 | 245.00 | 112.70 | 23.40 | 258.00 | 4.60 | 5.50 | 229.00 |
| | PO | PO06 | SC | 1 | 24.50 | 245.00 | 112.70 | 23.40 | 258.00 | 4.60 | 5.50 | 229.00 |
| | PO | PO07 | SO | 1 | 24.50 | 245.00 | 112.70 | 23.40 | 258.00 | 4.60 | 5.50 | 229.00 |
| | PO | PO09 | SO | 3 | 24.50 | 245.00 | 112.70 | 23.40 | 258.00 | 4.60 | 5.50 | 229.00 |
| | PO | PO11 | SA | 2 | 24.50 | 245.00 | 112.70 | 23.40 | 258.00 | 4.60 | 5.50 | 229.00 |
| | PO | PO13 | OP | 3 | 24.50 | 245.00 | 112.70 | 23.40 | 258.00 | 4.60 | 5.50 | 229.00 |
| | PO | PO14 | OP | 1 | 24.50 | 245.00 | 112.70 | 23.40 | 258.00 | 4.60 | 5.50 | 229.00 |
| | PO | PO15 | SO | 2 | 24.50 | 245.00 | 112.70 | 23.40 | 258.00 | 4.60 | 5.50 | 229.00 |
| | PO | PO17 | OP | 2 | 24.50 | 245.00 | 112.70 | 23.40 | 258.00 | 4.60 | 5.50 | 229.00 |
| | PO | PO19 | OP | 1 | 24.50 | 245.00 | 112.70 | 23.40 | 258.00 | 4.60 | 5.50 | 229.00 |
| | PO | PO23 | SO | 6 | 24.50 | 245.00 | 112.70 | 23.40 | 258.00 | 4.60 | 5.50 | 229.00 |
| | PO | PO24 | SO | 4 | 24.50 | 245.00 | 112.70 | 23.40 | 258.00 | 4.60 | 5.50 | 229.00 |
| | PO | PO25 | SO | 1 | 24.50 | 245.00 | 112.70 | 23.40 | 258.00 | 4.60 | 5.50 | 229.00 |
| | SR | SR18 | SC | 1 | 24.50 | 245.00 | 112.70 | 23.40 | 258.00 | 4.60 | 5.50 | 229.00 |
| | SR | SR21 | SC | 1 | 24.50 | 245.00 | 112.70 | 23.40 | 258.00 | 4.60 | 5.50 | 229.00 |
| | TE | TE04 | SC | 4 | 24.50 | 245.00 | 112.70 | 23.40 | 258.00 | 4.60 | 5.50 | 229.00 |

| | | | | | | | | | | | | | | |
|---|---|---|---|---|---|---|---|---|---|---|---|---|---|---|
| | | | TE | TE17 | SO | 1 | 24.50 | 245.00 | 112.70 | 23.40 | 258.00 | 4.60 | 5.50 | 229.00 |
| | | | TE | TE23 | SO | 2 | 24.50 | 245.00 | 112.70 | 23.40 | 258.00 | 4.60 | 5.50 | 229.00 |
| | | | TR | TR04 | SO | 1 | 24.50 | 245.00 | 112.70 | 23.40 | 258.00 | 4.60 | 5.50 | 229.00 |
| | | | TR | TR09 | OP | 2 | 24.50 | 245.00 | 112.70 | 23.40 | 258.00 | 4.60 | 5.50 | 229.00 |
| | | | TR | TR18 | SO | 3 | 24.50 | 245.00 | 112.70 | 23.40 | 258.00 | 4.60 | 5.50 | 229.00 |
| | | | TR | TR19 | SO | 4 | 24.50 | 245.00 | 112.70 | 23.40 | 258.00 | 4.60 | 5.50 | 229.00 |
| | | | TR | TR20 | SO | 3 | 24.50 | 245.00 | 112.70 | 23.40 | 258.00 | 4.60 | 5.50 | 229.00 |
| | | | VE | VE01 | OP | 1 | 24.50 | 245.00 | 112.70 | 23.40 | 258.00 | 4.60 | 5.50 | 229.00 |
| | | | VE | VE14 | SC | 1 | 24.50 | 245.00 | 112.70 | 23.40 | 258.00 | 4.60 | 5.50 | 229.00 |
| | | | VE | VE15 | SC | 2 | 24.50 | 245.00 | 112.70 | 23.40 | 258.00 | 4.60 | 5.50 | 229.00 |
| | | | VM | VM02 | SO | 1 | 24.50 | 245.00 | 112.70 | 23.40 | 258.00 | 4.60 | 5.50 | 229.00 |
| | | | VM | VM03 | SO | 1 | 24.50 | 245.00 | 112.70 | 23.40 | 258.00 | 4.60 | 5.50 | 229.00 |
| | | | VM | VM05 | SO | 1 | 24.50 | 245.00 | 112.70 | 23.40 | 258.00 | 4.60 | 5.50 | 229.00 |
| | | | VM | VM07 | SO | 2 | 24.50 | 245.00 | 112.70 | 23.40 | 258.00 | 4.60 | 5.50 | 229.00 |
| | | | VM | VM18 | OP | 1 | 24.50 | 245.00 | 112.70 | 23.40 | 258.00 | 4.60 | 5.50 | 229.00 |
| | | *Patagioenas plumbea* | BA | BA12 | SC | 1 | 23.10 | 340.00 | 141.60 | 23.10 | 316.47 | 4.30 | 5.10 | 178.80 |
| | | | PO | PO03 | SO | 1 | 23.10 | 340.00 | 141.60 | 23.10 | 316.47 | 4.30 | 5.10 | 178.80 |
| | | | TE | TE02 | SC | 1 | 23.10 | 340.00 | 141.60 | 23.10 | 316.47 | 4.30 | 5.10 | 178.80 |
| | | *Patagioenas subvinacea* | BA | BA18 | SC | 1 | 15.90 | 297.50 | 120.70 | 22.40 | 281.14 | 3.80 | 4.10 | 167.25 |
| | | | PO | PO11 | SO | 1 | 15.90 | 297.50 | 120.70 | 22.40 | 281.14 | 3.80 | 4.10 | 167.25 |
| | | | PO | PO19 | OP | 2 | 15.90 | 297.50 | 120.70 | 22.40 | 281.14 | 3.80 | 4.10 | 167.25 |
| | | | VM | VM04 | OP | 2 | 15.90 | 297.50 | 120.70 | 22.40 | 281.14 | 3.80 | 4.10 | 167.25 |
| | | *Zenaida auriculata* | TR | TR17 | OP | 2 | 17.50 | 250.00 | 78.95 | 19.00 | 200.00 | 3.70 | 4.00 | 110.10 |
| Passeriformes | Passerellidae | *Ammodramus aurifrons* | BA | BA12 | SC | 2 | 15.43 | 128.14 | 43.13 | 18.03 | 86.11 | 6.54 | 6.21 | 18.30 |
| | | | BA | BA18 | SC | 8 | 15.43 | 128.14 | 43.13 | 18.03 | 86.11 | 6.54 | 6.21 | 18.30 |
| | | | ES | ES02 | SC | 2 | 15.43 | 128.14 | 43.13 | 18.03 | 86.11 | 6.54 | 6.21 | 18.30 |
| | | | ES | ES12 | SC | 2 | 13.85 | 120.25 | 41.65 | 25.63 | 59.36 | 5.66 | 5.21 | 17.93 |
| | | | ES | ES14 | SC | 3 | 15.43 | 128.14 | 43.13 | 18.03 | 86.11 | 6.54 | 6.21 | 18.30 |
| | | | PO | PO13 | OP | 4 | 16.09 | 128.73 | 43.92 | 17.32 | 87.21 | 6.87 | 6.41 | 18.48 |
| | | | PO | PO14 | OP | 4 | 15.40 | 128.57 | 42.51 | 16.46 | 89.56 | 6.21 | 5.69 | 18.15 |
| | | | PO | PO17 | OP | 1 | 15.43 | 128.14 | 43.13 | 18.03 | 86.11 | 6.54 | 6.21 | 18.30 |
| | | | PO | PO18 | OP | 3 | 15.43 | 128.14 | 43.13 | 18.03 | 86.11 | 6.54 | 6.21 | 18.30 |
| | | | PO | PO22 | SC | 1 | 15.44 | 130.30 | 40.28 | 17.37 | 98.30 | 6.70 | 6.16 | 19.70 |

| | | | | | | | | | | | | |
|---|---|---|---|---|---|---|---|---|---|---|---|---|
| | PO | PO23 | SO | 1 | 15.43 | 128.14 | 43.13 | 18.03 | 86.11 | 6.54 | 6.21 | 18.30 |
| | PO | PO24 | SO | 2 | 15.43 | 128.14 | 43.13 | 18.03 | 86.11 | 6.54 | 6.21 | 18.30 |
| | SR | SR06 | SO | 2 | 15.43 | 128.14 | 43.13 | 18.03 | 86.11 | 6.54 | 6.21 | 18.30 |
| | TE | TE05 | SC | 3 | 15.43 | 128.14 | 43.13 | 18.03 | 86.11 | 6.54 | 6.21 | 18.30 |
| | TE | TE06 | SC | 2 | 15.43 | 128.14 | 43.13 | 18.03 | 86.11 | 6.54 | 6.21 | 18.30 |
| | TR | TR01 | OP | 2 | 15.43 | 128.14 | 43.13 | 18.03 | 86.11 | 6.54 | 6.21 | 18.30 |
| | TR | TR02 | OP | 2 | 15.43 | 128.14 | 43.13 | 18.03 | 86.11 | 6.54 | 6.21 | 18.30 |
| | TR | TR03 | OP | 2 | 15.43 | 128.14 | 43.13 | 18.03 | 86.11 | 6.54 | 6.21 | 18.30 |
| | TR | TR06 | SO | 2 | 15.43 | 128.14 | 43.13 | 18.03 | 86.11 | 6.54 | 6.21 | 18.30 |
| | TR | TR07 | SO | 2 | 15.43 | 128.14 | 43.13 | 18.03 | 86.11 | 6.54 | 6.21 | 18.30 |
| | TR | TR08 | OP | 2 | 15.43 | 128.14 | 43.13 | 18.03 | 86.11 | 6.54 | 6.21 | 18.30 |
| | TR | TR09 | OP | 3 | 15.43 | 128.14 | 43.13 | 18.03 | 86.11 | 6.54 | 6.21 | 18.30 |
| | TR | TR10 | OP | 2 | 15.43 | 128.14 | 43.13 | 18.03 | 86.11 | 6.54 | 6.21 | 18.30 |
| | TR | TR13 | OP | 2 | 15.43 | 128.14 | 43.13 | 18.03 | 86.11 | 6.54 | 6.21 | 18.30 |
| | TR | TR14 | OP | 3 | 15.43 | 128.14 | 43.13 | 18.03 | 86.11 | 6.54 | 6.21 | 18.30 |
| | TR | TR18 | SO | 1 | 22.24 | 127.65 | 47.98 | 18.93 | 90.95 | 11.72 | 11.09 | 19.10 |
| | TR | TR20 | SO | 3 | 15.43 | 128.14 | 43.13 | 18.03 | 86.11 | 6.54 | 6.21 | 18.30 |
| | TR | TR22 | OP | 1 | 15.43 | 128.14 | 43.13 | 18.03 | 86.11 | 6.54 | 6.21 | 18.30 |
| | TR | TR24 | OP | 2 | 15.43 | 128.14 | 43.13 | 18.03 | 86.11 | 6.54 | 6.21 | 18.30 |
| | TR | TR25 | OP | 2 | 15.43 | 128.14 | 43.13 | 18.03 | 86.11 | 6.54 | 6.21 | 18.30 |
| | VE | VE01 | OP | 1 | 15.43 | 128.14 | 43.13 | 18.03 | 86.11 | 6.54 | 6.21 | 18.30 |
| | VE | VE02 | OP | 3 | 15.58 | 131.84 | 43.58 | 17.51 | 89.42 | 6.53 | 5.93 | 17.65 |
| | VM | VM01 | SO | 2 | 15.43 | 128.14 | 43.13 | 18.03 | 86.11 | 6.54 | 6.21 | 18.30 |
| | VM | VM02 | SO | 2 | 15.43 | 128.14 | 43.13 | 18.03 | 86.11 | 6.54 | 6.21 | 18.30 |
| | VM | VM03 | SO | 3 | 15.43 | 128.14 | 43.13 | 18.03 | 86.11 | 6.54 | 6.21 | 18.30 |
| | VM | VM04 | OP | 3 | 14.00 | 128.72 | 42.99 | 16.87 | 88.71 | 5.75 | 6.00 | 18.24 |
| | VM | VM05 | SO | 1 | 15.43 | 128.14 | 43.13 | 18.03 | 86.11 | 6.54 | 6.21 | 18.30 |
| | VM | VM17 | OP | 1 | 13.28 | 129.00 | 42.91 | 16.28 | 90.00 | 5.34 | 5.88 | 18.20 |
| | VM | VM19 | OP | 2 | 15.43 | 128.14 | 43.13 | 18.03 | 86.11 | 6.54 | 6.21 | 18.30 |
| *Arremonops conirostris* | BA | BA07 | SO | 2 | 18.79 | 166.19 | 64.17 | 19.09 | 117.09 | 8.66 | 8.68 | 35.21 |
| | BA | BA08 | SO | 1 | 18.79 | 166.19 | 64.17 | 19.09 | 117.09 | 8.66 | 8.68 | 35.21 |
| | BA | BA09 | SO | 1 | 18.79 | 166.19 | 64.17 | 19.09 | 117.09 | 8.66 | 8.68 | 35.21 |
| | BA | BA10 | SC | 2 | 18.79 | 166.19 | 64.17 | 19.09 | 117.09 | 8.66 | 8.68 | 35.21 |

| | | | | | | | | | | | | | |
|---|---|---|---|---|---|---|---|---|---|---|---|---|---|
| | | BA | BA12 | SC | 3 | 18.79 | 166.19 | 64.17 | 19.09 | 117.09 | 8.66 | 8.68 | 35.21 |
| | | BA | BA13 | SC | 1 | 18.79 | 166.19 | 64.17 | 19.09 | 117.09 | 8.66 | 8.68 | 35.21 |
| | | BA | BA16 | OP | 4 | 18.79 | 166.19 | 64.17 | 19.09 | 117.09 | 8.66 | 8.68 | 35.21 |
| | | BA | BA18 | SC | 2 | 18.79 | 166.19 | 64.17 | 19.09 | 117.09 | 8.66 | 8.68 | 35.21 |
| | | PO | PO05 | SC | 2 | 18.79 | 166.19 | 64.17 | 19.09 | 117.09 | 8.66 | 8.68 | 35.21 |
| | | PO | PO14 | OP | 3 | 18.05 | 164.73 | 60.88 | 18.72 | 112.71 | 7.92 | 8.61 | 35.37 |
| | | PO | PO15 | SO | 1 | 18.79 | 166.19 | 64.17 | 19.09 | 117.09 | 8.66 | 8.68 | 35.21 |
| | | PO | PO18 | OP | 1 | 18.79 | 166.19 | 64.17 | 19.09 | 117.09 | 8.66 | 8.68 | 35.21 |
| | | PO | PO23 | SO | 6 | 18.79 | 166.19 | 64.17 | 19.09 | 117.09 | 8.66 | 8.68 | 35.21 |
| | | TE | TE05 | SC | 2 | 18.79 | 166.19 | 64.17 | 19.09 | 117.09 | 8.66 | 8.68 | 35.21 |
| | | TE | TE08 | SC | 2 | 18.79 | 166.19 | 64.17 | 19.09 | 117.09 | 8.66 | 8.68 | 35.21 |
| | | TE | TE14 | SC | 2 | 20.18 | 180.60 | 70.24 | 15.98 | 128.62 | 11.62 | 7.36 | 34.35 |
| | | TE | TE17 | SO | 2 | 18.79 | 166.19 | 64.17 | 19.09 | 117.09 | 8.66 | 8.68 | 35.21 |
| | | TR | TR01 | OP | 2 | 18.79 | 166.19 | 64.17 | 19.09 | 117.09 | 8.66 | 8.68 | 35.21 |
| | | TR | TR09 | OP | 2 | 18.79 | 166.19 | 64.17 | 19.09 | 117.09 | 8.66 | 8.68 | 35.21 |
| | | TR | TR15 | OP | 2 | 18.46 | 136.15 | 63.41 | 16.51 | 112.56 | 7.86 | 7.75 | 33.36 |
| | | TR | TR18 | SO | 4 | 20.54 | 167.63 | 62.50 | 20.79 | 112.30 | 9.25 | 9.82 | 36.11 |
| | | TR | TR20 | SO | 4 | 18.79 | 166.19 | 64.17 | 19.09 | 117.09 | 8.66 | 8.68 | 35.21 |
| | | TR | TR22 | OP | 2 | 18.79 | 166.19 | 64.17 | 19.09 | 117.09 | 8.66 | 8.68 | 35.21 |
| | | VM | VM07 | SO | 1 | 17.82 | 172.00 | 63.00 | 21.58 | 124.00 | 7.34 | 8.40 | 36.10 |
| | | VM | VM08 | SO | 1 | 18.52 | 172.70 | 63.70 | 22.28 | 124.70 | 8.04 | 9.10 | 36.80 |
| | | VM | VM17 | OP | 1 | 15.99 | 175.00 | 68.00 | 19.14 | 119.00 | 7.45 | 8.75 | 34.70 |
| | | VM | VM18 | OP | 1 | 15.94 | 174.95 | 67.95 | 19.09 | 118.95 | 7.40 | 8.70 | 34.65 |
| Thraupidae | *Sicalis flaveola* | BA | BA04 | OP | 2 | 16.51 | 145.00 | 54.41 | 13.53 | 123.51 | 6.07 | 7.71 | 21.80 |
| | | BA | BA05 | OP | 2 | 16.51 | 145.00 | 54.41 | 13.53 | 123.51 | 6.07 | 7.71 | 21.80 |
| | | BA | BA07 | SO | 1 | 16.51 | 145.00 | 54.41 | 13.53 | 123.51 | 6.07 | 7.71 | 21.80 |
| | | BA | BA09 | SO | 1 | 16.51 | 145.00 | 54.41 | 13.53 | 123.51 | 6.07 | 7.71 | 21.80 |
| | | BA | BA10 | SC | 2 | 16.51 | 145.00 | 54.41 | 13.53 | 123.51 | 6.07 | 7.71 | 21.80 |
| | | BA | BA12 | SC | 1 | 16.51 | 145.00 | 54.41 | 13.53 | 123.51 | 6.07 | 7.71 | 21.80 |
| | | BA | BA14 | SO | 2 | 16.51 | 145.00 | 54.41 | 13.53 | 123.51 | 6.07 | 7.71 | 21.80 |
| | | BA | BA15 | OP | 2 | 16.51 | 145.00 | 54.41 | 13.53 | 123.51 | 6.07 | 7.71 | 21.80 |
| | | BA | BA17 | SO | 6 | 16.51 | 145.00 | 54.41 | 13.53 | 123.51 | 6.07 | 7.71 | 21.80 |
| | | BA | BA18 | SC | 3 | 16.51 | 145.00 | 54.41 | 13.53 | 123.51 | 6.07 | 7.71 | 21.80 |

| | | | | | | | | | | | | |
|---|---|---|---|---|---|---|---|---|---|---|---|---|
| | PO | PO07 | SO | 2 | 16.51 | 145.00 | 54.41 | 13.53 | 123.51 | 6.07 | 7.71 | 21.80 |
| | PO | PO13 | OP | 2 | 16.51 | 145.00 | 54.41 | 13.53 | 123.51 | 6.07 | 7.71 | 21.80 |
| | PO | PO14 | OP | 2 | 16.51 | 145.00 | 54.41 | 13.53 | 123.51 | 6.07 | 7.71 | 21.80 |
| | PO | PO15 | SO | 2 | 16.51 | 145.00 | 54.41 | 13.53 | 123.51 | 6.07 | 7.71 | 21.80 |
| | PO | PO18 | OP | 3 | 16.51 | 145.00 | 54.41 | 13.53 | 123.51 | 6.07 | 7.71 | 21.80 |
| | PO | PO22 | SC | 3 | 16.51 | 145.00 | 54.41 | 13.53 | 123.51 | 6.07 | 7.71 | 21.80 |
| | PO | PO23 | SO | 2 | 16.51 | 145.00 | 54.41 | 13.53 | 123.51 | 6.07 | 7.71 | 21.80 |
| | PO | PO24 | SO | 3 | 16.51 | 145.00 | 54.41 | 13.53 | 123.51 | 6.07 | 7.71 | 21.80 |
| | PO | PO25 | SO | 2 | 16.51 | 145.00 | 54.41 | 13.53 | 123.51 | 6.07 | 7.71 | 21.80 |
| | TE | TE01 | SC | 3 | 16.51 | 145.00 | 54.41 | 13.53 | 123.51 | 6.07 | 7.71 | 21.80 |
| | TE | TE06 | SC | 2 | 16.51 | 145.00 | 54.41 | 13.53 | 123.51 | 6.07 | 7.71 | 21.80 |
| | TR | TR21 | OP | 2 | 16.51 | 145.00 | 54.41 | 13.53 | 123.51 | 6.07 | 7.71 | 21.80 |
| | VE | VE02 | OP | 2 | 16.51 | 145.00 | 54.41 | 13.53 | 123.51 | 6.07 | 7.71 | 21.80 |
| | VE | VE03 | SO | 2 | 16.51 | 145.00 | 54.41 | 13.53 | 123.51 | 6.07 | 7.71 | 21.80 |
| | VM | VM04 | OP | 3 | 16.51 | 145.00 | 54.41 | 13.53 | 123.51 | 6.07 | 7.71 | 21.80 |
| | VM | VM18 | OP | 2 | 16.51 | 145.00 | 54.41 | 13.53 | 123.51 | 6.07 | 7.71 | 21.80 |
| | VM | VM19 | OP | 2 | 16.51 | 145.00 | 54.41 | 13.53 | 123.51 | 6.07 | 7.71 | 21.80 |
| *Sporophila angolensis* | BA | BA07 | SO | 3 | 13.00 | 123.00 | 47.00 | 11.67 | 86.33 | 7.83 | 8.94 | 12.03 |
| | BA | BA08 | SO | 1 | 12.89 | 115.22 | 44.98 | 11.85 | 89.13 | 7.37 | 9.10 | 12.10 |
| | BA | BA14 | SO | 1 | 13.75 | 116.00 | 53.29 | 11.83 | 79.48 | 6.78 | 9.82 | 12.03 |
| | BA | BA17 | SO | 2 | 13.29 | 120.90 | 45.54 | 12.64 | 78.81 | 7.42 | 8.66 | 11.52 |
| | BA | BA18 | SC | 2 | 13.29 | 120.90 | 45.54 | 12.64 | 78.81 | 7.42 | 8.66 | 11.52 |
| | ES | ES03 | SC | 2 | 8.96 | 119.97 | 38.51 | 12.40 | 64.41 | 8.21 | 9.62 | 10.10 |
| | ES | ES06 | SC | 1 | 24.93 | 115.30 | 11.52 | 34.78 | 22.76 | 3.26 | 2.25 | 4.70 |
| | PO | PO03 | SO | 2 | 13.29 | 120.90 | 45.54 | 12.64 | 78.81 | 7.42 | 8.66 | 11.52 |
| | PO | PO11 | SO | 2 | 13.29 | 120.90 | 45.54 | 12.64 | 78.81 | 7.42 | 8.66 | 11.52 |
| | PO | PO18 | OP | 2 | 13.29 | 120.90 | 45.54 | 12.64 | 78.81 | 7.42 | 8.66 | 11.52 |
| | PO | PO24 | SO | 2 | 13.29 | 120.90 | 45.54 | 12.64 | 78.81 | 7.42 | 8.66 | 11.52 |
| | TE | TE06 | SC | 1 | 13.29 | 120.90 | 45.54 | 12.64 | 78.81 | 7.42 | 8.66 | 11.52 |
| | TE | TE14 | SC | 2 | 13.19 | 119.63 | 46.72 | 9.10 | 87.34 | 7.76 | 8.60 | 12.65 |
| | TR | TR18 | SO | 3 | 14.16 | 116.64 | 50.13 | 12.20 | 81.02 | 7.62 | 8.94 | 10.61 |
| | TR | TR19 | SO | 2 | 12.29 | 118.18 | 51.26 | 10.72 | 84.16 | 7.90 | 8.99 | 10.40 |
| | VE | VE02 | OP | 1 | 13.14 | 125.94 | 54.14 | 10.28 | 89.39 | 7.49 | 9.36 | 12.40 |

| | | | | | | | | | | | | |
|---|---|---|---|---|---|---|---|---|---|---|---|---|
| | VM | VM07 | SO | 1 | 12.76 | 128.00 | 54.05 | 11.26 | 86.00 | 7.96 | 9.27 | 11.80 |
| | VM | VM18 | OP | 1 | 12.21 | 137.00 | 42.00 | 10.98 | 80.20 | 6.90 | 8.35 | 19.20 |
| | TE | TE06 | SC | 2 | 11.00 | 100.00 | 37.50 | 13.80 | 91.33 | 4.60 | 5.70 | 7.80 |
| *Sporophila castaneiventris* | TE | TE21 | OP | 2 | 11.00 | 100.00 | 37.50 | 13.80 | 91.33 | 4.60 | 5.70 | 7.80 |
| | VM | VM01 | SO | 2 | 11.00 | 100.00 | 37.50 | 13.80 | 91.33 | 4.60 | 5.70 | 7.80 |
| *Sporophila crassirostris* | VE | VE02 | OP | 2 | 13.56 | 122.38 | 49.90 | 10.86 | 89.79 | 7.60 | 8.74 | 11.60 |
| | VE | VE19 | SC | 1 | 12.87 | 111.19 | 41.39 | 10.93 | 75.55 | 8.67 | 8.29 | 11.00 |
| *Sporophila intermedia* | TR | TR18 | SO | 1 | 12.31 | 104.21 | 37.19 | 5.57 | 76.55 | 8.04 | 8.89 | 12.20 |
| *Sporophila minuta* | TR | TR18 | SO | 1 | 9.15 | 90.00 | 38.99 | 14.15 | 86.85 | 5.00 | 6.20 | 7.90 |
| | BA | BA07 | SO | 4 | 26.38 | 125.00 | 48.48 | 11.37 | 90.50 | 7.63 | 8.11 | 14.75 |
| *Sporophila murallae* | BA | BA13 | SC | 2 | 26.38 | 125.00 | 48.48 | 11.37 | 90.50 | 7.63 | 8.11 | 14.75 |
| | TR | TR18 | SO | 2 | 26.38 | 125.00 | 48.48 | 11.37 | 90.50 | 7.63 | 8.11 | 14.75 |
| *Sporophila nigricollis* | PO | PO23 | SO | 2 | 9.39 | 111.00 | 45.00 | 20.00 | 94.30 | 6.04 | 5.84 | 9.80 |
| | BA | BA07 | SO | 3 | 10.95 | 107.50 | 39.79 | 10.57 | 72.10 | 6.19 | 6.33 | 9.60 |
| | BA | BA10 | SC | 1 | 13.08 | 102.24 | 39.12 | 11.01 | 74.31 | 9.20 | 6.84 | 9.20 |
| | BA | BA12 | SC | 1 | 13.08 | 102.24 | 39.12 | 11.01 | 74.31 | 9.20 | 6.84 | 9.20 |
| | BA | BA14 | SO | 2 | 13.08 | 102.24 | 39.12 | 11.01 | 74.31 | 9.20 | 6.84 | 9.20 |
| | BA | BA16 | OP | 2 | 13.08 | 102.24 | 39.12 | 11.01 | 74.31 | 9.20 | 6.84 | 9.20 |
| | BA | BA17 | SO | 8 | 13.08 | 102.24 | 39.12 | 11.01 | 74.31 | 9.20 | 6.84 | 9.20 |
| | ES | ES07 | SC | 1 | 13.08 | 102.24 | 39.12 | 11.01 | 74.31 | 9.20 | 6.84 | 9.20 |
| | ES | ES14 | SC | 2 | 13.08 | 102.24 | 39.12 | 11.01 | 74.31 | 9.20 | 6.84 | 9.20 |
| | PO | PO03 | SO | 2 | 13.08 | 102.24 | 39.12 | 11.01 | 74.31 | 9.20 | 6.84 | 9.20 |
| | PO | PO09 | SO | 2 | 13.08 | 102.24 | 39.12 | 11.01 | 74.31 | 9.20 | 6.84 | 9.20 |
| *Volatinia jacarina* | PO | PO11 | SO | 4 | 13.08 | 102.24 | 39.12 | 11.01 | 74.31 | 9.20 | 6.84 | 9.20 |
| | PO | PO14 | OP | 2 | 13.08 | 102.24 | 39.12 | 11.01 | 74.31 | 9.20 | 6.84 | 9.20 |
| | PO | PO15 | SO | 2 | 13.08 | 102.24 | 39.12 | 11.01 | 74.31 | 9.20 | 6.84 | 9.20 |
| | PO | PO16 | SC | 2 | 13.08 | 102.24 | 39.12 | 11.01 | 74.31 | 9.20 | 6.84 | 9.20 |
| | PO | PO17 | OP | 1 | 13.08 | 102.24 | 39.12 | 11.01 | 74.31 | 9.20 | 6.84 | 9.20 |
| | PO | PO19 | OP | 2 | 13.08 | 102.24 | 39.12 | 11.01 | 74.31 | 9.20 | 6.84 | 9.20 |
| | PO | PO22 | SC | 2 | 13.08 | 102.24 | 39.12 | 11.01 | 74.31 | 9.20 | 6.84 | 9.20 |
| | PO | PO23 | SO | 2 | 13.08 | 102.24 | 39.12 | 11.01 | 74.31 | 9.20 | 6.84 | 9.20 |
| | PO | PO24 | SO | 4 | 13.08 | 102.24 | 39.12 | 11.01 | 74.31 | 9.20 | 6.84 | 9.20 |
| | PO | PO25 | SO | 2 | 13.08 | 102.24 | 39.12 | 11.01 | 74.31 | 9.20 | 6.84 | 9.20 |

| | | | | | | | | | | | | | | |
|---|---|---|---|---|---|---|---|---|---|---|---|---|---|---|
| | | | SR | SR02 | SO | 1 | 10.72 | 103.00 | 43.69 | 10.71 | 79.46 | 6.44 | 6.76 | 10.30 |
| | | | TE | TE06 | SC | 1 | 13.08 | 102.24 | 39.12 | 11.01 | 74.31 | 9.20 | 6.84 | 9.20 |
| | | | TE | TE21 | OP | 1 | 13.08 | 102.24 | 39.12 | 11.01 | 74.31 | 9.20 | 6.84 | 9.20 |
| | | | TR | TR02 | OP | 2 | 13.08 | 102.24 | 39.12 | 11.01 | 74.31 | 9.20 | 6.84 | 9.20 |
| | | | TR | TR03 | OP | 2 | 13.08 | 102.24 | 39.12 | 11.01 | 74.31 | 9.20 | 6.84 | 9.20 |
| | | | TR | TR07 | SO | 1 | 13.08 | 102.24 | 39.12 | 11.01 | 74.31 | 9.20 | 6.84 | 9.20 |
| | | | TR | TR08 | OP | 2 | 13.08 | 102.24 | 39.12 | 11.01 | 74.31 | 9.20 | 6.84 | 9.20 |
| | | | TR | TR09 | OP | 1 | 13.08 | 102.24 | 39.12 | 11.01 | 74.31 | 9.20 | 6.84 | 9.20 |
| | | | TR | TR10 | OP | 3 | 13.08 | 102.24 | 39.12 | 11.01 | 74.31 | 9.20 | 6.84 | 9.20 |
| | | | TR | TR13 | OP | 6 | 13.08 | 102.24 | 39.12 | 11.01 | 74.31 | 9.20 | 6.84 | 9.20 |
| | | | TR | TR14 | OP | 2 | 13.08 | 102.24 | 39.12 | 11.01 | 74.31 | 9.20 | 6.84 | 9.20 |
| | | | TR | TR18 | SO | 4 | 15.03 | 99.24 | 38.99 | 11.14 | 75.07 | 9.09 | 7.69 | 8.40 |
| | | | TR | TR19 | SO | 1 | 13.81 | 89.00 | 34.10 | 10.75 | 73.22 | 6.19 | 6.58 | 9.10 |
| | | | TR | TR22 | OP | 2 | 13.08 | 102.24 | 39.12 | 11.01 | 74.31 | 9.20 | 6.84 | 9.20 |
| | | | TR | TR24 | OP | 2 | 13.08 | 102.24 | 39.12 | 11.01 | 74.31 | 9.20 | 6.84 | 9.20 |
| | | | TR | TR25 | OP | 2 | 13.08 | 102.24 | 39.12 | 11.01 | 74.31 | 9.20 | 6.84 | 9.20 |
| | | | VE | VE01 | OP | 1 | 13.08 | 102.24 | 39.12 | 11.01 | 74.31 | 9.20 | 6.84 | 9.20 |
| | | | VE | VE02 | OP | 2 | 13.08 | 102.24 | 39.12 | 11.01 | 74.31 | 9.20 | 6.84 | 9.20 |
| | | | VE | VE03 | SO | 2 | 13.08 | 102.24 | 39.12 | 11.01 | 74.31 | 9.20 | 6.84 | 9.20 |
| | | | VM | VM01 | SO | 3 | 13.08 | 102.24 | 39.12 | 11.01 | 74.31 | 9.20 | 6.84 | 9.20 |
| | | | VM | VM03 | SO | 2 | 13.08 | 102.24 | 39.12 | 11.01 | 74.31 | 9.20 | 6.84 | 9.20 |
| | | | VM | VM04 | OP | 3 | 13.08 | 102.24 | 39.12 | 11.01 | 74.31 | 9.20 | 6.84 | 9.20 |
| | | | VM | VM17 | OP | 1 | 13.30 | 111.00 | 38.11 | 12.41 | 73.85 | 6.18 | 5.33 | 10.20 |
| | | | VM | VM18 | OP | 2 | 13.08 | 102.24 | 39.12 | 11.01 | 74.31 | 9.20 | 6.84 | 9.20 |
| Tinamiformes | Tinamidae | *Crypturellus cinereus* | SR | SR05 | SC | 1 | 32.40 | 305.00 | 64.45 | 49.95 | 237.50 | 5.20 | 5.40 | 541.05 |
| | | | SR | SR15 | SC | 2 | 32.40 | 305.00 | 64.45 | 49.95 | 237.50 | 5.20 | 5.40 | 541.05 |
| | | | TE | TE08 | SC | 1 | 32.40 | 305.00 | 64.45 | 49.95 | 237.50 | 5.20 | 5.40 | 541.05 |
| | | *Crypturellus soui* | BA | BA12 | SC | 1 | 21.35 | 225.00 | 46.45 | 38.20 | 200.00 | 4.20 | 3.90 | 218.10 |
| | | | PO | PO03 | SO | 2 | 21.35 | 225.00 | 46.45 | 38.20 | 200.00 | 4.20 | 3.90 | 218.10 |
| | | | PO | PO05 | SC | 1 | 21.35 | 225.00 | 46.45 | 38.20 | 200.00 | 4.20 | 3.90 | 218.10 |
| | | | PO | PO13 | OP | 1 | 21.35 | 225.00 | 46.45 | 38.20 | 200.00 | 4.20 | 3.90 | 218.10 |
| | | | VE | VE14 | SC | 1 | 21.35 | 225.00 | 46.45 | 38.20 | 200.00 | 4.20 | 3.90 | 218.10 |
| | | *Crypturellus undulatus* | BA | BA12 | SC | 1 | 34.00 | 220.00 | 56.60 | 48.00 | 160.00 | 6.00 | 5.80 | 564.40 |

| | | | | | | | | | | | | |
|---|---|---|---|---|---|---|---|---|---|---|---|---|
| | PO | PO03 | SO | 1 | 34.00 | 220.00 | 56.60 | 48.00 | 160.00 | 6.00 | 5.80 | 564.40 |
| | PO | PO12 | SO | 1 | 34.00 | 220.00 | 56.60 | 48.00 | 160.00 | 6.00 | 5.80 | 564.40 |
| | PO | PO13 | OP | 1 | 34.00 | 220.00 | 56.60 | 48.00 | 160.00 | 6.00 | 5.80 | 564.40 |
| | SR | SR06 | SO | 1 | 34.00 | 220.00 | 56.60 | 48.00 | 160.00 | 6.00 | 5.80 | 564.40 |
| | TE | TE12 | SC | 1 | 34.00 | 220.00 | 56.60 | 48.00 | 160.00 | 6.00 | 5.80 | 564.40 |
| | VE | VE23 | SC | 2 | 34.00 | 220.00 | 56.60 | 48.00 | 160.00 | 6.00 | 5.80 | 564.40 |
| | BA | BA12 | SC | 2 | 22.60 | 120.00 | 61.80 | 37.20 | 180.26 | 5.00 | 5.30 | 352.10 |
| *Tinamus guttatus* | BA | BA18 | SC | 1 | 22.60 | 120.00 | 61.80 | 37.20 | 180.26 | 5.00 | 5.30 | 352.10 |
| | SR | SR15 | SC | 1 | 22.60 | 120.00 | 61.80 | 37.20 | 180.26 | 5.00 | 5.30 | 352.10 |

BA: Batalla 13; PO: El Porvenir; VE: La Vega; TE: El Tesoro; TR: El Triunfo; ES: La Esmeralda; SR: Santa Rosa; VM: Villa Mery; TC: tree cover; OP: open; SO: semi-open; SC: semi-closed; AB: abundance of individuals; ALT: bill height; COM: commissure; AEX: extended wing; LTA: tarsus length; LCO: tail length; LTO: total body length; CTO: total culmen; PES: body weight (g).

**Table S3. Matrix of granivorous bird species assigned to two functional groups (small and large) based on the dendrogram generated by hierarchical cluster analysis using the ward.D algorithm.**

| # | FG | Order | Familiy | Scientific name | AB | Morphological functional traits | | | | | | | |
|---|---|---|---|---|---|---|---|---|---|---|---|---|---|
| | | | | | | **CTO** | **LTO** | **LCO** | **LTA** | **AEX** | **COM** | **ALT** | **PES** |
| 01 | FG_S | Passeriformes | Passerellidae | *Ammodramus aurifrons* | 89 | 15.49 | 128.14 | 43.15 | 18.08 | 86.23 | 6.60 | 6.28 | 18.33 |
| 02 | | | | *Arremonops conirostris* | 57 | 18.68 | 166.75 | 64.34 | 19.20 | 117.49 | 8.60 | 8.69 | 35.23 |
| 03 | | | Thraupidae | *Sicalis flaveola* | 61 | 16.51 | 145.00 | 54.41 | 13.53 | 123.51 | 6.07 | 7.71 | 21.80 |
| 04 | | | | *Sporophila angolensis* | 31 | 13.60 | 120.94 | 45.39 | 13.04 | 78.05 | 7.30 | 8.57 | 11.54 |
| 05 | | | | *Sporophila castaneiventris* | 06 | 11.00 | 100.00 | 37.50 | 13.80 | 91.33 | 4.60 | 5.70 | 7.80 |
| 06 | | | | *Sporophila crassirostris* | 03 | 13.22 | 116.79 | 45.65 | 10.90 | 82.67 | 8.14 | 8.52 | 11.30 |
| 07 | | | | *Sporophila intermedia* | 01 | 12.31 | 104.21 | 37.19 | 5.57 | 76.55 | 8.04 | 8.89 | 12.20 |
| 08 | | | | *Sporophila minuta* | 01 | 9.15 | 90.00 | 38.99 | 14.15 | 86.85 | 5.00 | 6.20 | 7.90 |
| 09 | | | | *Sporophila murallae* | 08 | 26.38 | 125.00 | 48.48 | 11.37 | 90.50 | 7.63 | 8.11 | 14.75 |
| 10 | | | | *Sporophila nigricollis* | 02 | 9.39 | 111.00 | 45.00 | 20.00 | 94.30 | 6.04 | 5.84 | 9.80 |
| 11 | | | | *Volatinia jacarina* | 96 | 13.09 | 102.14 | 39.10 | 11.02 | 74.37 | 8.93 | 6.83 | 9.22 |
| 12 | | Columbiformes | Columbidae | *Columbina minuta* | 15 | 16.03 | 148.08 | 55.22 | 11.75 | 122.63 | 5.99 | 3.43 | 35.47 |
| 13 | | | | *Columbina talpacoti* | 49 | 18.08 | 157.57 | 65.00 | 12.69 | 133.85 | 4.54 | 4.43 | 43.06 |
| | | | | | **M** | **15.21** | **128.68** | **47.63** | **14.87** | **93.75** | **7.51** | **7.31** | **18.65** |
| | | | | | **SD** | **2.64** | **22.99** | **9.67** | **3.80** | **20.58** | **1.45** | **1.18** | **9.05** |
| 14 | FG_L | Tinamiformes | Tinamidae | *Crypturellus soui* | 04 | 32.40 | 305.00 | 64.45 | 49.95 | 237.50 | 5.20 | 5.40 | 541.05 |
| 15 | | | | *Crypturellus cinereus* | 06 | 21.35 | 225.00 | 46.45 | 38.20 | 200.00 | 4.20 | 3.90 | 218.10 |
| 16 | | | | *Crypturellus undulatus* | 08 | 34.00 | 220.00 | 56.60 | 48.00 | 160.00 | 6.00 | 5.80 | 564.40 |
| 17 | | | | *Tinamus guttatus* | 04 | 22.60 | 220.00 | 71.80 | 37.20 | 180.26 | 5.00 | 5.30 | 352.10 |
| 18 | | Columbiformes | Columbidae | *Leptotila rufaxilla* | 14 | 27.37 | 240.33 | 89.20 | 22.76 | 187.33 | 6.21 | 5.08 | 119.27 |
| 19 | | | | *Patagioenas cayennensis* | 94 | 24.50 | 245.00 | 112.70 | 23.40 | 258.00 | 4.60 | 5.50 | 229.00 |
| 20 | | | | *Patagioenas plumbea* | 03 | 23.10 | 340.00 | 141.60 | 23.10 | 316.47 | 4.30 | 5.10 | 178.80 |
| 21 | | | | *Patagioenas subvinacea* | 06 | 15.90 | 297.50 | 120.70 | 22.40 | 281.14 | 3.80 | 4.10 | 167.25 |
| 22 | | | | *Zenaida auriculata* | 02 | 17.50 | 250.00 | 78.95 | 19.00 | 200.00 | 3.70 | 4.00 | 110.10 |
| | | | | | **M** | **23.28** | **223.02** | **89.31** | **24.15** | **210.63** | **4.88** | **4.99** | **203.69** |
| | | | | | **SD** | **4.88** | **50.14** | **27.19** | **10.40** | **57.65** | **0.92** | **0.70** | **143.13** |

FG: functional group; GF_S: functional group of small granivorous birds; FG_L: functional group of large granivorous birds; AB: abundance of individuals; ALT: bill height; COM: commissure; AEX: extended wing; LTA: tarsus length; LCO: tail length; LTO: total body length; CTO: total culmen; PES: body weight (g); M: trait mean; SD: standard deviation.

**Table S4. Linear Models Analysis and pairwise Fisher's mean comparisons (α = 0.05) of the functional traits of granivorous bird species across three types of tree cover in livestock landscapes of the Colombian Amazon.**

| Traits | Tree cover | Mean | SE | N | Group | F | p-value |
|---|---|---|---|---|---|---|---|
| CTO | SC | 19.83 | 0.62 | 72 | ***A*** | 6.69 | 0.0015 |
| | SO | 18.57 | 0.50 | 113 | ***A*** | | |
| | OP | 16.79 | 0.57 | 87 | ***B*** | | |
| LTO | SC | 176.72 | 6.79 | 72 | ***A*** | 3.82 | 0.0232 |
| | SO | 165.87 | 5.42 | 113 | ***AB*** | | |
| | OP | 151.64 | 6.18 | 87 | ***B*** | | |
| LTA | SC | 21.61 | 0.97 | 72 | ***A*** | 8.47 | 0.0003 |
| | SO | 17.61 | 0.77 | 113 | ***B*** | | |
| | OP | 16.45 | 0.88 | 87 | ***B*** | | |
| AEX | SC | 149.69 | 8.05 | 72 | ***A*** | 3.56 | 0.0297 |
| | SO | 142.16 | 6.42 | 113 | ***A*** | | |
| | OP | 122.3 | 7.32 | 87 | ***B*** | | |
| PES | SC | 127.76 | 14.60 | 72 | ***A*** | 6.52 | 0.0017 |
| | SO | 87.59 | 11.60 | 113 | ***B*** | | |
| | OP | 56.64 | 13.30 | 87 | ***B*** | | |

ALT: bill height; COM: commissure; AEX: extended wing; LTA: tarsus length; LCO: tail length; LTO: total body length; CTO: total culmen; PES: body weight (g); SO: semi-open; OP: open; SC: semi-closed; SE: standard error. Values sharing the same letter indicate means that are not significantly different (p > 0.05).

**Table S5. Linear Models Analysis and pairwise Fisher's mean comparisons (α = 0.05) of functional diversity indices of granivorous birds across three types of tree cover in livestock landscapes of the Colombian Amazon.**

| Functional diversity index | Tree cover | Mean | Group | F | p-value |
|---|---|---|---|---|---|
| FRic | OP | 0.136 | ***A*** | 1.38 | 0.2633 |
| | SO | 0.320 | ***A*** | | |
| | SC | 0.602 | ***A*** | | |
| FEve | OP | 0.420 | ***A*** | 1.14 | 0.3316 |
| | SO | 0.458 | ***A*** | | |
| | SC | 0.667 | ***A*** | | |
| FDiv | OP | 0.718 | ***A*** | 0.99 | 0.3794 |
| | SO | 0.743 | ***A*** | | |
| | SC | 0.736 | ***A*** | | |
| FDis | OP | 1.811 | ***AB*** | 7.12 | 0.0013 |
| | SO | 2.134 | ***A*** | | |
| | SC | 2.123 | ***B*** | | |
| RaoQ | OP | 4.123 | ***B*** | 7.41 | 0.0001 |
| | SO | 5.038 | ***A*** | | |
| | SC | 5.085 | ***B*** | | |

FRic: functional richness; FEve: functional evenness; FDiv: functional divergence; FDis: functional dispersion; RaoQ: Rao's quadratic entropy; OP: open; SO: semi-open; SC: semi-closed. Values sharing the same letter indicate means that are not significantly different ($p > 0.05$).
